# Inherited maternal memory T cells protect offspring from respiratory viral infection through disease tolerance

**DOI:** 10.64898/2026.10.30.763208

**Authors:** Hamadoun Touré, Brittany Jimena, Madelyn E. Cabán, Emma E. Lederer, Diane M. Rico, Addison Medlin McFadden, Saajidah Abideen, Pranav Kokati, Whitney E. Harrington, Meghan A. Koch

## Abstract

Maternal immune transfer is a powerful mechanism for early-life protection from infection, yet the full repertoire of protective factors remains unknown. We discovered that maternal memory T cells are both necessary and sufficient to protect offspring from lethal respiratory infection. This antigen-specific protection operated independently of maternal antibodies and offspring T cell responses and required IL-10 produced by maternal T cells. Transfer of protective maternal T cells reduced offspring inflammatory responses and lung tissue damage without affecting viral burdens, indicating that maternal memory T cells directly suppress pathogenic innate inflammatory responses to preserve offspring health. Critically, maternal vaccination similarly protected offspring in a T cell-dependent manner. These findings reveal maternal antigen-specific T cell transfer as a complementary immunological defense mechanism to protect offspring against pathogens that escape maternal antibodies, with broad implications for maternal vaccination strategies.

**Graphical abstract:** 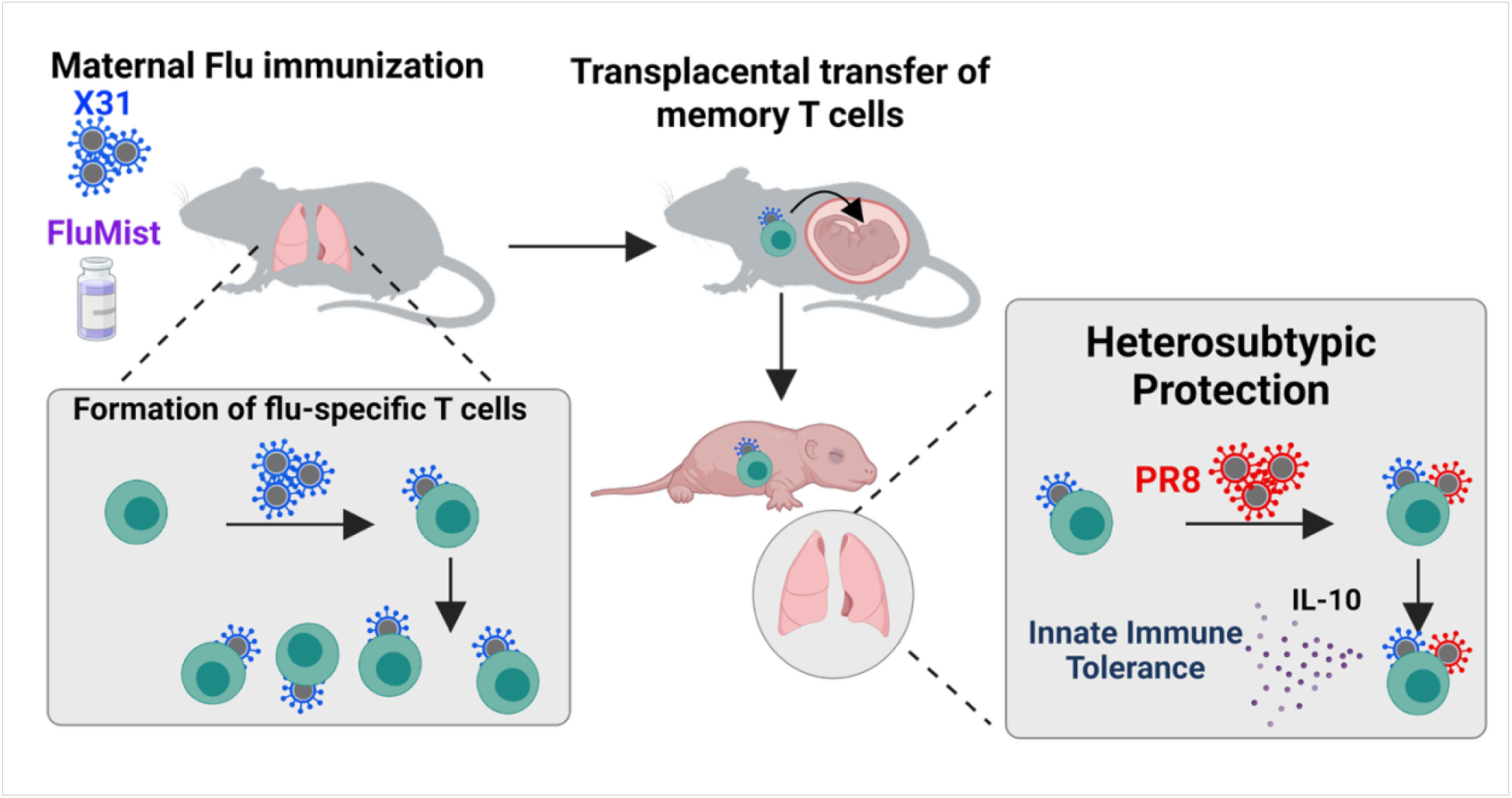

## Introduction

The mammalian neonate must tolerate innocuous antigens following exposure to milk, the environment, and commensal microbes after birth. To achieve this, multiple mechanisms drive preferential regulatory responses in early life, including immature antigen-presenting cells, a skewed T cell receptor (TCR) repertoire, increased frequency of regulatory T cells, and altered B cell responses^1,2^. However, these processes also increase the risk of neonatal susceptibility to infection, including to invasive bacterial and respiratory tract infections, which could lead to a large evolutionary penalty^3–6^. Maternal immune transfer partially counters this penalty. The passive transmission of soluble maternal complement proteins or cytokines can enhance offspring immunity to enteric infections in an antigen-agnostic manner^7,8^. Further, transfer of maternal IgG across the placenta and IgA via breastfeeding provide offspring well-characterized antigen-specific defense, offering inherited protection directly matched to the maternal immunological experience. However, this protection is limited: maternal antibody titers wane within months, and clinically relevant pathogens, including influenza A virus (IAV), HIV, and SARS-CoV-2, routinely mutate surface antigens to evade antibody recognition^9–11^. When antigenic escape occurs, maternal antibody-mediated protection collapses, potentially leaving infants exposed^11^.

Aside from soluble immune proteins, maternal cells are vertically transmitted to and engraft within offspring, a process termed ‘maternal microchimerism’^12^. Recent evidence shows that this maternal graft is enriched for antigen-experienced memory T cells^13,14^, a central arm of the adaptive immune response that mediates long-lived immune defense. Unlike antibodies and other soluble immune factors, memory T cells can survive many years and self-renew, even in the absence of cognate antigen^15–17^. Additionally, memory T cells can recognize conserved, pathogen-derived epitopes distinct from those targeted by antibodies, providing complementary, durable defense against pathogens that evolve to evade antibody-mediated neutralization^18^. These observations raised a key question: can maternal memory T cells defend offspring from infection when antibodies fail? If so, what mechanisms underlie this protection?

Here we report that maternal antigen-specific T cells are necessary and sufficient to protect offspring from lethal respiratory infection. We show that protective maternal T cells produce IL-10 to regulate innate immunity and prevent lung immunopathology in offspring, and that maternal vaccination can similarly elicit this mode of transgenerational protection.

## Results

### Pre-conception maternal IAV immunity protects offspring from heterosubtypic infection

To evaluate whether maternal T cells protect offspring from infection, we used a well-established mouse model of heterosubtypic IAV infection ^5,19,20,20–23^. In this system, memory T cells generated in response to sublethal infection of a host with one strain of IAV (X31; H3N2) are necessary and sufficient to protect the host against a subsequent lethal challenge with a heterosubtypic strain (PR8; H1N1)^21,20^. Critically, neutralizing antibody responses to IAV are strain-specific; antibodies elicited by one strain do not protect against infection with the heterosubtypic strain^24–26^. This feature of the model thus allowed us to directly assess maternal T cell function in immunocompetent mothers without confounding effects from cross-reactive maternal antibodies. We intranasally (i.n.) infected C57BL/6NTac dams from Taconic with a sublethal dose of PR8, resulting in weight loss between days 4 and 7 post-infection (p.i.) and a recovery phase starting at day 8 p.i. **(Fig S1A).** At day 10 p.i., after viral clearance and generation of IAV nucleoprotein (NP)-specific T cells in the lungs **(Fig S1B, C)**, we paired PR8 immune dams (PR8_pre_) with uninfected sires. We separately paired control, uninfected dams with uninfected sires. We then challenged age-matched offspring of both types of breeding pairs with a lethal dose of heterosubtypic X31 (30PFU) between postnatal days 3-5 (p3-5) and monitored their survival (**Fig 1A**). Whereas 58% of offspring of naïve dams succumbed to infection, mortality was significantly reduced to 5% in pups of PR8 preconception-infected (PR8_pre_) dams (**Fig 1B**). Complementary experiments using X31 immune dams (X31_pre_ - **Fig S1A-C)** revealed a similar survival benefit in pups infected with heterosubtypic PR8 (**Fig 1C**). Offspring of IAV-immune dams also exhibited reduced morbidity, evidenced by greater weight gain (**Fig S1D-E**).

**Figure 1:**
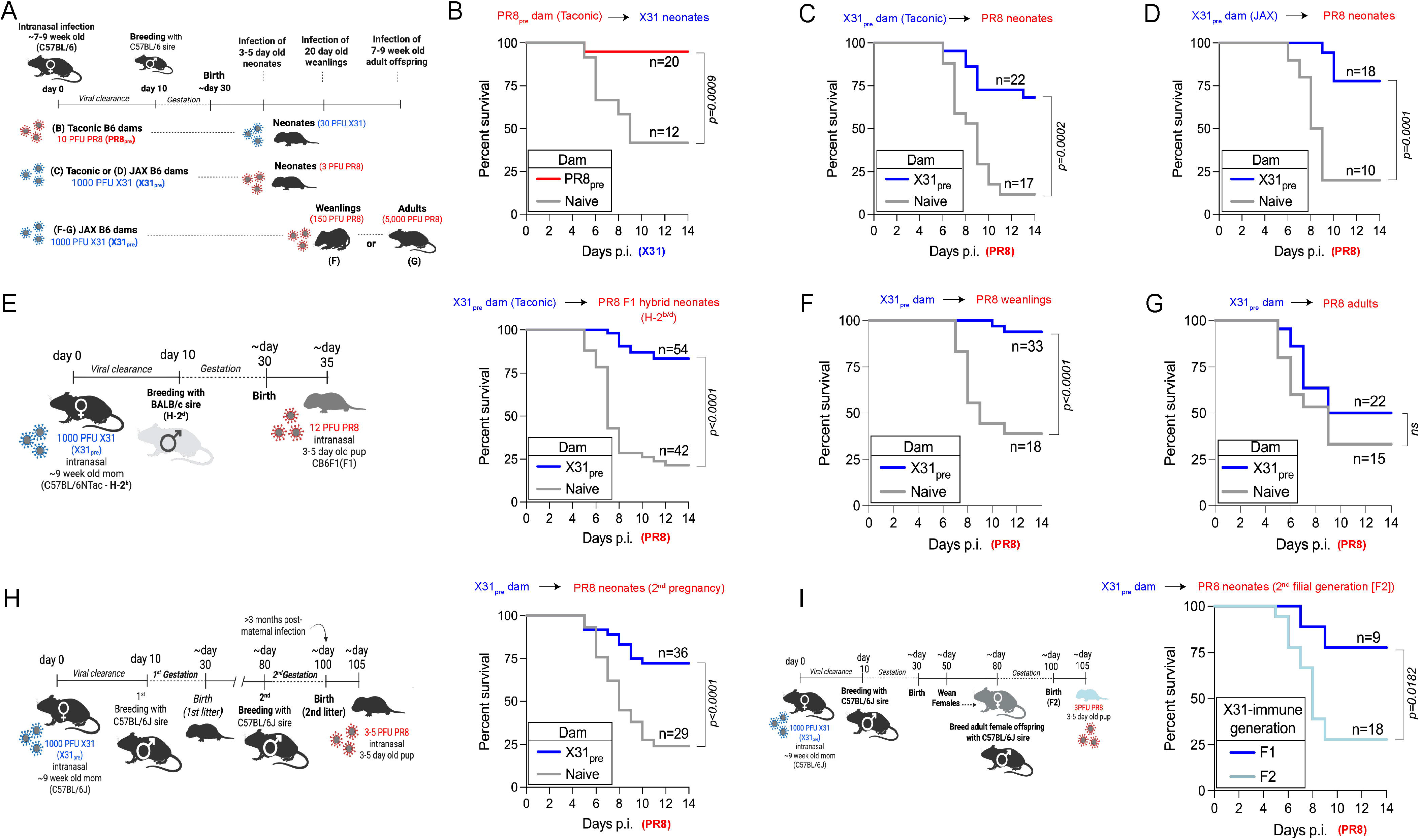
Preconception maternal IAV infection confers heterosubtypic protection to offspring. **(A)** Experimental approach of B-D and F-G. **(B)** Survival of pups born to naïve or PR8-immune (PR8_pre_) C57BL/6NTac dams and infected with X31 between day 3-5 post birth. **(C-D)** Survival of pups born to naïve or X31-immune (X31_pre_) C57BL/6NTac dams (C) or C57BL/6J dams (D) and infected with PR8 between day 3-5 post birth. **(E)** Experimental approach (left panel) and survival (right panel) of pups born to naïve or X31_pre_ C57BL/6NTac dams crossed to BALB/c sires (F1). Pups were infected with PR8 between day 3-5 post birth. **(F)** Pooled survival of males and females weanling born to naïve or X31_pre_ C57BL/6J dams and infected with PR8 between day 18-20 post birth. **(G)** Pooled survival of males and females adult born to naïve or X31_pre_ C57BL/6J dams and infected with PR8 between weeks 7-9 post birth. **(H)** Experimental approach (left panel) and survival (right panel) of second litter pups born to naïve or X31_pre_ C57BL/6J dams crossed to C57BL/6J sires. Pups were infected with PR8 between day 3-5 post birth. **(I)** Experimental approach (upper panel) and survival (lower panel) of the first filial progeny (F1) or second filial progeny (F2) born from X31_pre_ dams. Pups were infected with PR8 between day 3-5 post birth. Statistical significance for survival curves was determined by the log-rank test (Mantel-Cox).

As host genetics and microbiota composition can substantially influence immune responses^27–29^, we repeated these experiments using X31_pre_ C57BL/6J dams sourced from Jackson Laboratory, which harbor a distinct microbiota composition and minor genetic variations compared to C57BL/6NTac mice^30–32^. Offspring maintained a significant survival advantage upon PR8 challenge (**Fig 1D**), establishing that transgenerational protection is a generalizable phenomenon. We further tested whether protection extends to MHC half-matched offspring, which better represents natural outbreeding and human genetic diversity. We generated F1 H-2^b/d^ MHC heterozygous offspring by crossing C57BL/6NTac dams (H-2^b^) with naïve BALB/c sires (H-2^d^) and challenged them with PR8 (12 PFU; dose titration shown in **Fig S1F**). F1 hybrid offspring of X31_pre_ dams exhibited significantly reduced mortality compared to those from naïve dams (**Fig 1E**). Collectively, these findings demonstrate that preconception maternal immunization with IAV confers robust heterosubtypic protection to both genetically identical and diverse (haplo-identical) offspring, suggesting a broadly protective mechanism across diverse immunogenetic contexts.

We next assessed the durability of this form of maternal immune defense by challenging older offspring born to naïve and X31_pre_ dams. Protection was evident in weaned male and female offspring separated from their mother and infected at p21, a developmental period akin to late infancy/early adolescence in humans^33,34^ (**Fig 1F; S2A)**. These findings indicate that protection is not simply mediated by re-stimulation of maternal immunity through contact with infected offspring. Notably, maternal immune status did not impact infection outcomes of adult offspring challenged with PR8 between 7-9 weeks of age (**Fig 1G and S2B**), indicating that this form of transgenerational immune defense operates primarily during early life. To determine whether this protection persisted beyond offspring conceived immediately after the acute maternal infection period, we challenged a second litter of offspring of X31_pre_ dams born more than three months after initial maternal exposure. These later-born offspring remained significantly protected from lethal PR8 challenge **(Fig 1H)**. Although IAV-derived antigens have been reported to persist for several weeks after pathogen clearance^35^, the extended interval between maternal infection and offspring protection argues against the transfer of residual IAV antigens from immune dams to pups as a mechanism of protection. We next tested whether this transgenerational protection extends to subsequent generations. While the first generation of offspring (F1) of X31_pre_ dams were protected, second filial generation (F2) pups remained highly susceptible to PR8 infection (**Fig 1I**), indicating that maternal-mediated heterosubtypic protection is restricted to a single generation. Collectively, these findings reveal a mode of transgenerational immune defense wherein pre-conceptional maternal immunity to respiratory viral infection mediates early-life offspring defense from related viral strains across multiple litters.

### Maternal antibodies and serum soluble factors are dispensable for transgenerational immune protection from heterosubtypic IAV infection

Maternal soluble and cellular factors involved in shaping offspring immunity can be transferred to offspring during gestation and via breastfeeding^36^. To distinguish the developmental period in which the protective maternal-derived benefit was transferred, we cross-fostered pups born to naïve or X31_pre_ dams at birth (**Fig. 2A**). In this setting, offspring born to X31_pre_ dams and nursed by naïve dams receive protective factors from immune dams during gestation but not through breastmilk; and vice versa. Offspring born to X31_pre_ dams remained protected from PR8 infection regardless of nursing mother. In contrast, pups born to naïve dams remained highly susceptible to infection even when nursed by X31_pre_ dams. Thus, heterosubtypic protection of young (p3-5) offspring is gestationally transmitted from dam to offspring.

**Figure 2:**
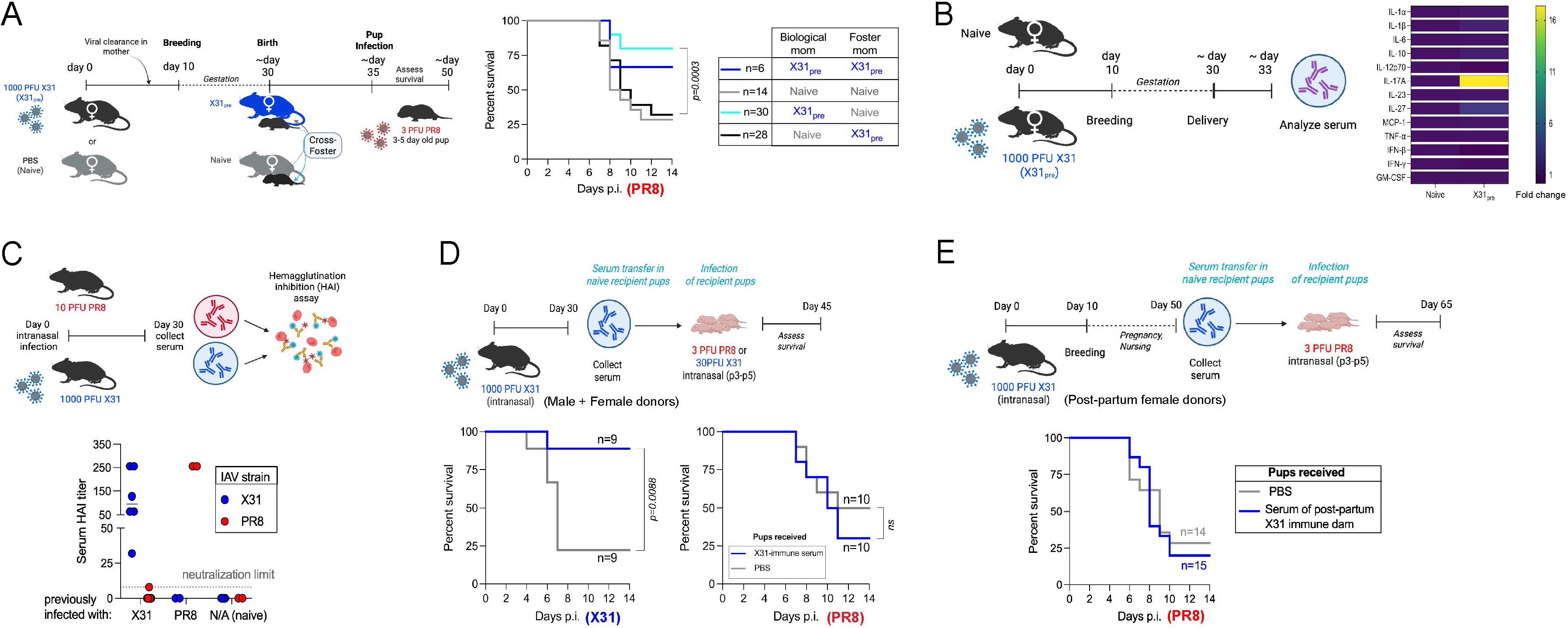
Maternal antibodies and serum soluble factors are dispensable for transgenerational immune protection from heterosubtypic IAV infection. (A) Experimental approach (left panel) and survival (right panel) of offspring born to and nursed by naïve (grey) or X31_pre_ (blue) dams or cross-fostered at birth (cyan and black). Pups were infected with PR8 between day 3-5 post birth. (B) Heatmap of serum cytokines levels fold change in post-partum X31_pre_ dams relative to post-partum naïve dams. (C) Quantification of serum hemagglutination inhibition (HAI) titer of naïve and infected adult mice on day 30 post-infection (p.i.). The HAI was determined as the reciprocal of the last serum dilution to inhibit hemagglutination. N/A: non-infected; neutralization limit = 8. (D) Upper panel: Experimental approach showing serum collection from X31-immune mice at day 30 p.i. and serum adoptive transfer in naïve recipient pups on the same day. Lower panel: p3-5 recipient were infected with X31 (left) or PR8 (right) and survival was monitored. (E) Upper panel: Experimental approach showing serum collection from post-partum X31_pre_ dams and serum adoptive transfer in naïve recipient pups on the same day. Lower panel: p3-5 recipient were infected with X31 and survival was monitored. Statistical significance for survival curves was determined by the log-rank test (Mantel-Cox).

The gestational transfer of maternal cytokines such as IL-6 can bias mucosal T cell responses, resulting in increased host defense to enteric infection^8^. However, serum cytokines were largely similar between naïve and X31_pre_ dams at day 3 post-partum, except for increased serum IL-17A and IL-27 in X31_pre_ dams **(Figure 2B and S3A)**.

Transplacental transfer of pathogen-specific antibodies represents a robust mechanism of maternal immune defense^37^. However, PR8 and X31 heterosubtypic strains lack cross-reactive neutralizing epitopes due to divergent hemagglutinin sequences^24–26^, which we confirmed via hemagglutination inhibition (HAI) assays using serum from X31- and PR8-immunized dams collected 30 days post-infection (**Fig 2C**). Importantly, antibodies can mediate host protection beyond neutralization via additional effector mechanisms including opsonization, complement activation, antibody-dependent cellular cytotoxicity, and antibody-dependent cellular phagocytosis^38^. While maternal antibody-mediated protection has been shown to occur in a strain-specific manner^39^, we tested whether non-neutralizing antibody effector functions could mediate heterosubtypic protection by transferring sera from X31-immune donors into naïve recipient pups. X31-immune serum significantly protected offspring from homotypic X31 infection but failed to protect against heterosubtypic PR8 challenge (**Fig 2D**). Likewise, sera from PR8 immune donors protected offspring from PR8 infection (**Fig S3B**), confirming that antibody-mediated protection is IAV strain-specific. Recent work has shown that pregnancy can induce post-translational modifications of maternal antibodies, enhancing their protective functions in offspring^40^. As an additional control, we transferred sera from postpartum X31_pre_ dams into naïve recipient pups and found that pups exhibited mortality following PR8 infection comparable to PBS controls (**Fig 2E**), excluding both conventional and pregnancy-modified antibodies as mediators of transgenerational protection. Collectively, these findings indicate that transmission of maternal defense against heterosubtypic infection is mediated via gestation through a mechanism that extends beyond maternal antibodies and other serum soluble immune factors.

### Maternal T cells confer transgenerational heterosubtypic defense against IAV

Studies in both humans and murine models have demonstrated that maternal T cells can be transmitted to offspring^41–43,13,44–47^. Additionally, rare cases of human infants with severe combined immunodeficiency have documented expanded populations of cytomegalovirus- or Epstein-Barr virus-specific maternal T cells following infection, indicating that maternal T cells can provide compensatory immune defense in offspring^48,49^. Given these observations, we hypothesized that maternal T cells confer heterosubtypic immune defense against influenza.

To directly test the functional relevance of transferred maternal T cells in conferring heterosubtypic protection, we developed an approach to specifically ablate inherited maternal T cells in offspring using CD4.Cre x ROSA^lslDTR^ (T^DTR^) dams. In these mice, all T cells express the diphtheria toxin receptor (DTR) and can be specifically depleted by administration of diphtheria toxin (DT)^50^ ^51^ (**Fig S4A,B**). To establish the utility of this approach for specifically manipulating inherited maternal T cells, we adoptively transferred total splenocytes from T^DTR^ mice harboring T cells that also express zsGreen1 (*i.e*., CD4Cre^+^xROSAl^slDTR/WT^ x ROSA^lslzsG/WT^ donors, referred to as T^DTR-zsG^) into p4-p7 congenically distinct recipient pups. DT treatment efficiently depleted donor T cells in pups, while sparing other donor populations (such as B cells) and recipient host-derived (i.e., pup-derived) T cells (**Fig S4C,D**).

We next infected Cre+ hemizygous T^DTR^ mothers with X31 and bred them 10 days post-infection. As control groups, we also bred naive T^DTR^ and X31_pre_ wild-type dams with B6 sires. We then cross-fostered Cre-offspring to lactating naïve dams at p5, treated them with DT or PBS, and infected them with PR8 (**Fig. 3B**). In this approach, Cre-pups do not express DTR, and thus only maternal DTR+ T cells can be depleted by DT. Additionally, although pups are capable of inheriting maternal T cells both *in utero* and via breastfeeding for the first 5 days of life, cross-fostering offspring to naïve lactating dams at p5 precludes the further potential transfer of maternal IAV-elicited T cells. Strikingly, DT treatment of Cre-, PR8-challenged offspring of T^DTR^ dams abrogated the protective benefit of preconception maternal X31 infection, resulting in increased mortality and morbidity (**Fig. 3C**). These effects were specific to maternal T cell depletion, as DT treatment did not impair survival of offspring from X31_pre_ wild-type (non-DTR) dams (**Fig 3C**), ruling out off-target DT toxicity.

**Figure 3:**
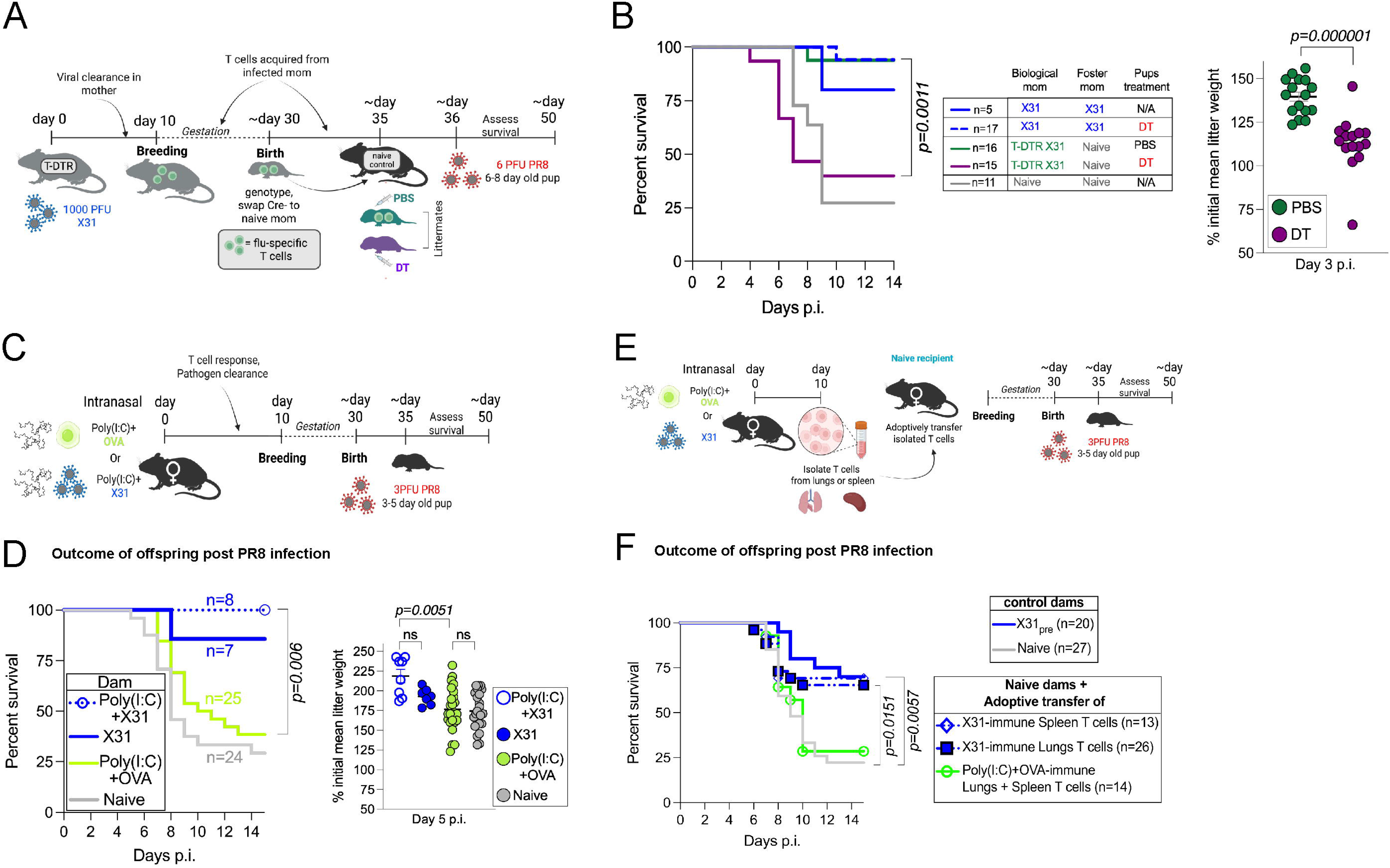
Maternal IAV-specific T cells are necessary and sufficient to confer transgenerational heterosubtypic protection. **(A)** Experimental approach to test whether maternal-derived, IAV-specific T cells (represented in green), confer transgenerational protection. **(B)** Left panel: Survival of littermate offspring born from X31-immune T^DTR^ dams, reared by naïve B6 dams, and given PBS (green) or DT (purple). T^DTR^: CD4.Cre^+/-^ x iDTR^lslFlox/wt^. Right panel: Percentage of initial body weight at d5 p.i. relative to the mean litter weight at d0. **(C)** Experimental approach to test whether maternal Poly(I:C)+OVA immunization protects offspring against PR8 infection. **(D)** Survival of pups born to naïve, Poly(I:C)+OVA or Poly(I:C)+X31-immune dams and infected with PR8 between day 3-5 post birth. **(E)** Experimental approach to test whether IAV-specific T cells are sufficient to protect offspring against PR8 infection. **(F)** Survival of pups born from control (naïve or X31_pre_) or naïve dams that received adoptive transfer of T cells from the lungs or spleen of X31-immune dams or pooled T cells from the lungs and spleen of Poly(I:C)+OVA-immune dams. Pups were infected with PR8 between day 3-5 post birth.

These findings support a direct effect of IAV-antigen-specific maternal T cells in mediating transgenerational protection. However, emerging studies indicate that maternal T cells elicited by vaccination or infection can enhance neonatal immunity in an antigen-agnostic manner, and preconception maternal immunization has been reported to augment transmission of maternal cells ^52,53,43,45,54^. To determine whether transgenerational protection depends on antigen-specific maternal T cells, we i.n. immunized dams with OVA protein adjuvanted with the double stranded RNA TLR3 agonist poly(I:C), modeling viral infection^55,56^. This strategy elicited robust CD4 and CD8 (including OVA-specific) lung T_RM_ cells, as well as effector memory (T_EM_), and central memory (T_CM_) populations in the lungs and the spleen (**Fig 3C and Fig S5A,B**). Offspring of OVA/polyi:c immunized dams were susceptible to PR8 infection, whereas offspring of X31_pre_ dams or X31-immune dams also given i.n. poly(I:C) remained protected (**Fig 3D**). In complementary experiments, we infected dams i.n. with *Listeria monocytogenes* expressing chromosomally integrated OVA (Lm-OVA). Like X31, this intracellular pathogen induces strong type-1 inflammatory responses, and i.n. infection elicited robust accumulation of pathogen-specific lung T_RM_ and T_EM_ as well as systemic effector populations^57,58^ (**Fig S5B-E**). Female mice recovered normally from Lm-OVA infection, with complete bacterial clearance by day 10 p.i. (**Fig S5C**). Consistent with our results using OVA/polyi:c immunized mothers, offspring of Lm-OVA-immune dams remained highly susceptible to PR8 challenge, whereas control offspring of X31_pre_ dams were protected (**Fig S5F**). Collectively, these data establish that transgenerational immune defense requires IAV-specific maternal T cells and cannot be recapitulated by non-specific maternal T cell activation, supporting an antigen-specific mechanism.

### Maternal IAV-elicited T cells are sufficient to confer transgenerational heterosubtypic protection to healthy and immunodeficient offspring

We next assessed whether maternal T cells from IAV immune mothers were sufficient to mediate heterosubtypic protection of offspring. In adult mice, lung-resident memory T cells (T_RM_) confer superior protection against heterosubtypic IAV infection compared to circulating memory populations^19,20^. To determine whether this paradigm extends to the maternal-offspring dyad, we transferred either lung-derived T cells (5×10⁵ CD4+ and 10×10⁵ CD8+ cells, predominantly T_RM_ and T_EM_; **Fig S5G**) or splenic T cells (5×10⁶ CD4+ and 10×10⁶ CD8+, predominantly T_CM_ and T_EM_; **Fig S5G**) from X31-immune donors into naïve recipient dams, then mated recipients with sires and challenged offspring with PR8 (**Fig 3E**). Strikingly, offspring of both groups were similarly protected, comparable to offspring of X31_pre_ dams, whereas control offspring born to naïve, non-transferred dams succumbed to infection (**Fig 3F**). As a key control, offspring of naïve dams given 5×10⁶ CD4⁺ and 5×10⁶ CD8⁺ T cells from OVA/poly(I:C)–challenged mice were not protected from PR8 infection (**Fig 3F**). Taken together, these data show that maternal T cells from IAV-immune mothers are sufficient to confer heterosubtypic protection to offspring independent of tissue origin, suggesting redundancy between resident and circulating memory T cell compartments in mediating transgenerational defense.

### Maternal memory T cells promote innate disease tolerance to protect offspring from lethal IAV

Host survival following infection, including IAV, depends on two complementary defense mechanisms: *(i)* resistance, which limits pathogen abundance, and *(ii)* disease tolerance, which minimizes infection-induced immunopathology without necessarily impacting pathogen loads^59,60^. Memory T cell-mediated protection against secondary heterosubtypic IAV infection has been shown to involve both mechanisms^20,22^. Offspring of X31_pre_ dams exhibited significantly reduced overall lung cellular infiltration and tissue damage at days 3 and 5 p.i. compared to offspring of naïve dams (**Fig 4A,B and Fig S6A**). However, similar viral loads were observed between groups early following infection (day 3 p.i.), and at day 5p.i., a timepoint immediately prior to mortality in susceptible pups (**Fig 4C**). This dissociation between pathogen control and survival suggests that maternal T cells primarily operate by mitigating lung immunopathology and promoting disease tolerance.

**Figure 4:**
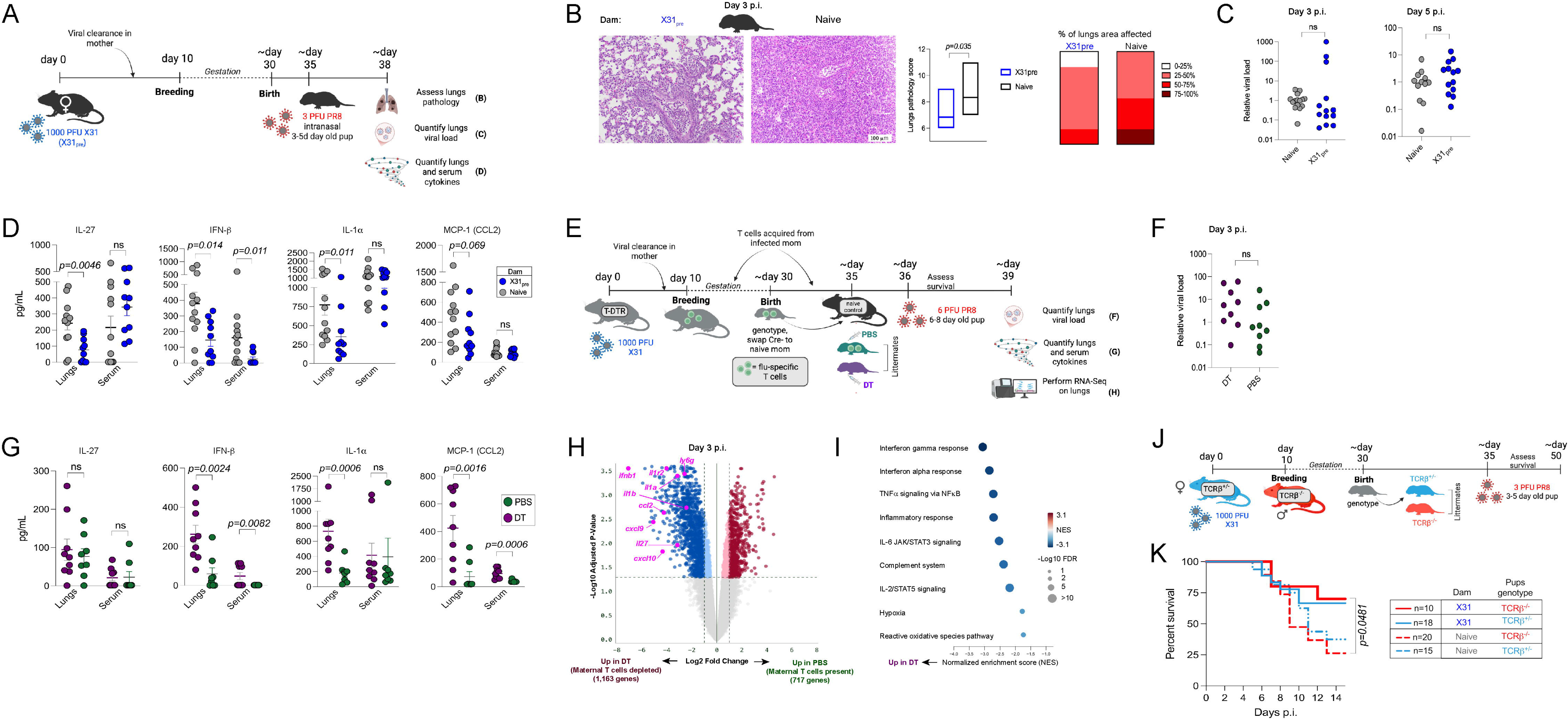
Maternal memory T cells confer transgenerational heterosubtypic protection by promoting disease tolerance. **(A)** Experimental approach to profile lungs disease and immune responses in offspring born from naïve and X31_pre_ dams. **(B)** Upper panel: Representative lungs H&E staining at day 3 post PR8 infection of pups born from naïve (right) or X31_pre_ dams (left). Lower panel: Cumulative lungs pathology score (left) and percentage of lungs area affected by the pathology (right); n=6 mice per group. **(C)** Relative PR8 viral load in the lungs of pups born from naïve or X31_pre_ dams. Viral *NS1* transcripts levels were normalized to β-*actin*, and the relative viral load were determined in function of the mean titer of the naïve condition either at day 3 (left) or day 5p.i. (right). **(D)** Cytokine levels (day 3 p.i.) in the lungs and serum of pups born from naïve or X31_pre_ dams and infected with PR8. **(E)** Experimental approach to profile lungs disease and immune responses in littermate offspring born from X31-immune T^DTR^, reared by naïve B6 dams, and treated with PBS or DT, then infected with PR8. **(F)** Relative PR8 viral load (day 3 p.i.) in the lungs of littermate pups born from X31-immune T^DTR^ and treated with PBS or DT. Viral *NS1* transcripts levels were normalized to β-*actin*, and the relative viral load were determined in function of the mean of the PBS-treated condition either at day 3 p.i. **(G)** Cytokine levels (day 3 p.i.) in the lungs and serum of littermate pups born from X31-immune T^DTR^ and treated with PBS or DT, then infected with PR8. **(H)** Volcano plot of differentially expressed genes at day 3 post-PR8 infection in the lungs of littermate pups born from X31-immune T^DTR^ and treated with PBS or DT. **(I)** Functional enrichment analysis showing the significant pathways in which the genes enriched in the DT-treated pups are involved. **(J)** Experimental approach to test whether maternal IAV immunity protects T cell deficient offspring against PR8 infection. **(K)** Survival of TCRβ^+/-^ and TCRβ^-/-^ offspring born from naïve or X31-immune TCRβ^+/-^ dams. Pups were infected with PR8 between day 3-5 post birth. Each dot represents an individual pup in B-E. Statistical significance was determined by the log-non-parametric Student’s t-test (Mann-Whitney) in B-E.

To further explore this idea, we profiled inflammatory cytokine and chemokine levels in the lungs and serum of infected offspring of naïve or X31_pre_ dams. Antiviral cytokine levels, including IL-6, IL-23, TNF-α, and IFN-γ, were equivalently high in both groups (**Fig. S6B**). However, offspring of naïve dams harbored significantly elevated levels of additional inflammatory factors including MCP-1, IL-27, IL-1-α, and IFN-β in the lungs at day 3 p.i. **(Fig 4D**), with levels converging by day 5 p.i. (**Fig S6C**). In the absence of offspring infection, comparable low levels of MCP-1, IL-27, IL-1α/β, and IFN-α were observed (**Fig S6D–E**), establishing that differences in infection-elicited cytokines were not driven by inherited ‘baseline’ cytokine levels.

Previous studies have demonstrated that maternal immune status and prior immune experience can influence immune predisposition in offspring^8,61,53,62^. Targeted transcriptomic analysis (NanoString) revealed modest differences in the lungs of uninfected X31_pre_ offspring compared to those born to naïve controls—including elevated levels of *Ebi2, Nod2, Cxcr3, Cxcr5, Il7r, tnfsf14, Ccl3*, and *Pdcd1lg2* (**Fig S6E**). To directly establish that maternal T cells regulate neonatal immunity to heterosubtypic infection and exclude potential maternal confounders related to prior immune experience, we compared littermate Cre-offspring of X31-immune T^DTR^ dams. These pups were either treated with PBS (maternal T cells retained) or DT (maternal T cells depleted) (**Fig 4E**). We observed similar pathogen burden in PBS- and DT-treated littermates at day 3 p.i. (**Fig 4F**), confirming that maternal T cells do not alter viral loads in offspring. However, largely recapitulating the dysregulation seen in offspring of naïve dams (**Fig 4C**), we found that maternal T cell-depleted pups harbored elevated MCP-1, IL-1α, and IFN-β (but not IL-27) at day 3 p.i. (**Fig 4G**). RNA-seq analysis revealed an enrichment of lung transcripts encoding for IFN-β (top hit), CCL2 (MCP-1), IL-1α/β, and IL-27 in DT-treated, maternal T cell depleted pups at day 3 p.i. (**Fig 4H**). Pathway enrichment analysis revealed that transcripts enriched in the lungs of DT-treated pups are primarily involved in inflammation, including responses to type I and II interferons, reactive oxidative species, and hypoxia (**Fig 4I**). Taken together, these findings indicate that maternal IAV-specific memory T cells protect offspring by orchestrating a coordinated innate immune response that mitigates immunopathology and early immune dysregulation.

The mitigation of early innate inflammatory responses by maternal IAV-specific T cells support a model in which maternal T cells primarily regulate offspring innate immune responses. To evaluate whether maternal T cells are sufficient to protect offspring even in the absence of pup-derived T cells, we infected TCRβ^+/-^ dams (which possess functional T cells) with X31 and subsequently mated them with TCRβ^-/-^ sires lacking αβ T cells. Progeny from this breeding produced both TCRβ^+/-^ and TCRβ^-/-^ that received maternal flu-specific memory T cells **(Fig 4J)**. Following PR8 infection, both TCRαβ+ T cell-sufficient and -deficient offspring were protected compared to age and genotype-matched control pups born to naïve dams **(Fig 4K)**. These findings were supported by complementary experiments in which the T cell repertoire of progeny was largely restricted to recognize ovalbumin (OVA), an irrelevant antigen not expressed by influenza virus. Again, both CD8 (OT-I) and CD4 (OT-II) TCR restricted and wild-type (non-transgenic) progeny born to X31_pre_ dams were protected against PR8 infection (**Fig S7A-D**, respectively), indicating that maternal influenza-specific T cells are sufficient to provide transgenerational defense against heterosubtypic infection even when the offspring T cell compartment is compromised.

### Maternal memory T cells primarily engage the offspring innate immune cells

To better characterize the mechanism of transgenerational heterosubtypic protection by maternal memory T cells, we firstly characterize their phenotypic features and the cells they engage in offspring lungs following infection. We confirmed that maternal T cells are transmitted to offspring and localize to the lungs following IAV infection. To do so, we generated CD4.Cre^+^ x ROSA^lslzsGreen1^ mice (herein referred to as T^zsG^) in which all T cells express the pH-stable fluorescent protein zsGreen1 **(Fig S8A)**. We next infected T^zsG^ dams with X31, bred them with wild-type sires at day 10 p.i., and infected resulting Cre-offspring with PR8. We detected maternal zsGreen1+ T cells in the offspring lungs at day 3 p.i., including nucleoprotein NP366-372 tetramer+ IAV-specific CD8 T cells^63^ **(Fig 5A)**. The majority of the maternal cells exhibited an effector memory phenotype and were not marked by intravenously injected anti-CD45, establishing that these maternal-derived cells reside in the offspring lung tissues and do not localize into the vasculature ^64^ (**Fig S8B**). As an important note, maternal T cells were extremely rare in offspring, and their detection was not consistent across individual pups and between litters using flow cytometric analysis, rendering their extensive characterization challenging. However, *zsGreen1*-speciifc mRNA transcripts were consistently detected in individual pups using a sensitive molecular approach, Countable PCR (**Fig 5B**), suggesting that live maternal T cells are present in offspring lungs following PR8 infection. These results further indicate that flow cytometric analysis of enzymatically digested lungs is not an optimal approach to track maternal T cells, given their low numbers.

**Figure 5:**
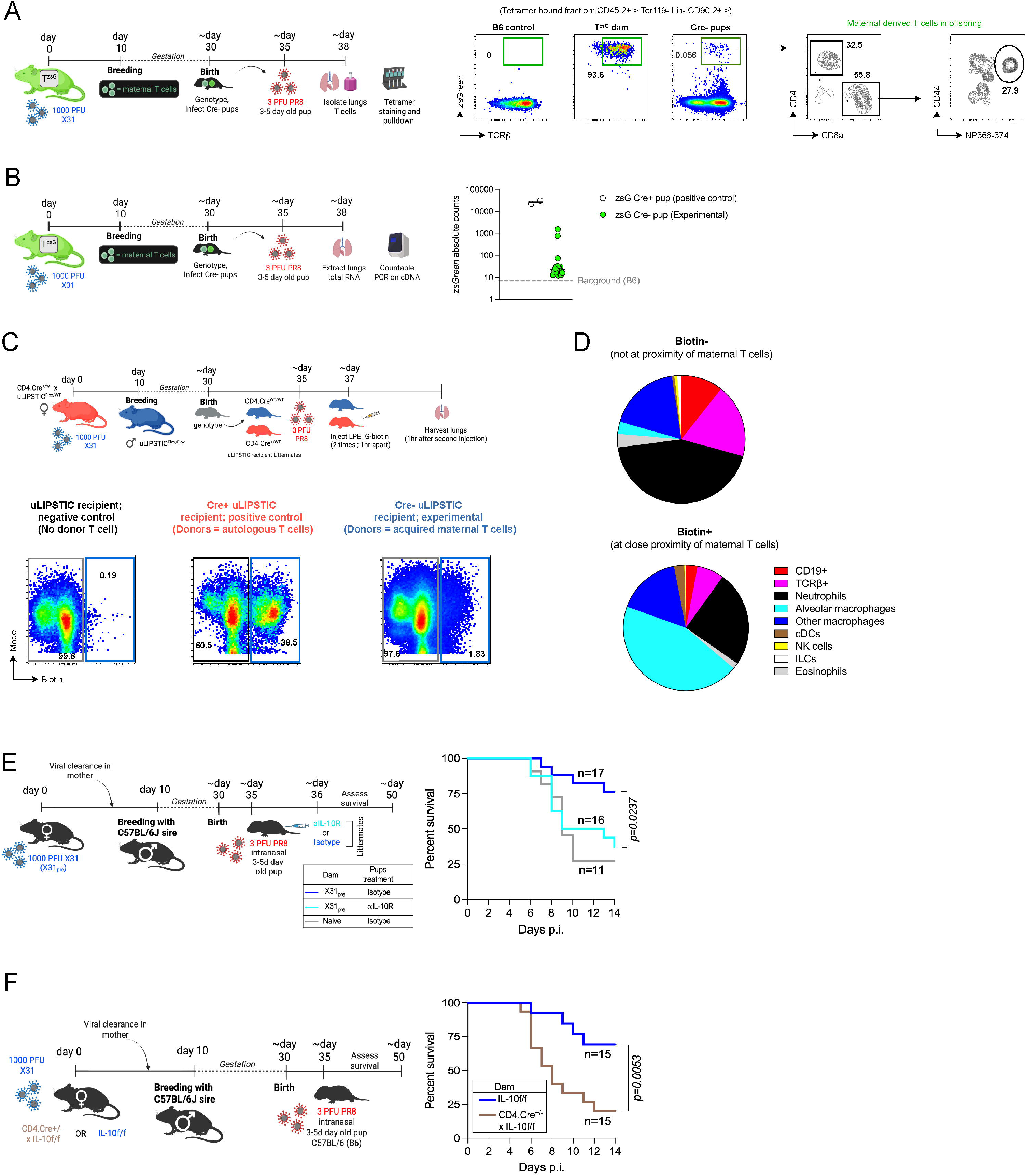
Maternal T cells engage the offspring innate immune cells and confer IAV heterosubtypic protection in an IL-10-dependent manner. **(A)** Experimental approach (left panel) and flow cytometric analysis of maternal microchimeric T cells (zsGreen+) in the offspring lungs following IAV infection (right panel). Lungs T cells were isolated, stained with tetramer, and the tetramer-enriched fraction was analyzed by flow cytometry. T^zsG^: CD4.Cre^+/-^ x zsGreen1^lslFlox/wt^; Lin: B220, CD19, Siglec-F, Ly6G. **(B)** Experimental approach (left panel) and detection of maternal microchimeric T cells (zsGreen transcripts) by Countable PCR in the offspring lungs following IAV infection (right panel). Total RNA were extracted from the lungs and reverse transcribed into cDNA. Countable PCR targeting *zsGreen*-specific cDNA was performed. **(C)** Upper panel: experimental approach to track cellular partner of maternal T cells in the lungs of the offspring following PR8 infection. Lower panel: Representative flow plots of the biotin signal in the lungs of the indicated recipient pups at 2 days post-PR8 infection. Pre-gate: CD45+ Ter119-. **(D)** Pie chart showing the proportions of different immune cell type proportions in the biotin- and biotin+ lung fractions of the experimental pups (Cre-recipients in which maternal T cells are donors). **(E)** Experimental approach (left) and survival (right) of littermate pups born X31_pre_ dams and treated with αIL-10R or isotype control at day 1 post-PR8 infection. **(F)** Experimental approach (left) and survival (right) of pups born from X31-immune IL-10flox/flox (IL-10f/f) or CD4.Cre^+/-^ x IL-10flox/flox dams. Each dot represents an individual pup in G. Statistical significance for survival curves was determined by the log-rank test (Mantel-Cox).

To directly identify offspring cells interacting with maternal memory T cells, we leveraged the universal LIPSTIC (uLIPSTIC) system, which enables recording of cell-cell interactions in vivo ^65^. We infected CD4.Cre^+/WT^ x uLIPSTIC^f/WT^ (referred as donor dams) with X31, and paired them with uLIPSTIC^f/f^ sires at day 10 p.i. The resulting Cre-negative offspring (recipients) were subsequently infected with PR8 and injected systemically with LPETG-biotin at day 2 p.i. In this setting, maternal donor T cells (transferred during gestation) label any cell of the recipient pups encountered in close proximity (<40nm) by transmitting the LPETG-biotin substrate. The interacting cells can then be detected by biotin staining **(Fig 5C)**. Biotin+ cells were composed predominantly of alveolar (44%) and other macrophage populations (16%), and neutrophils (25%), identifying these innate populations as major cellular partners of maternal T cells in the infected offspring lung **(Fig 5D)**.

### Maternal memory T cells promote disease tolerance in an IL-10 dependent manner

One mechanism by which memory T cells promote disease tolerance to protect the host from secondary heterosubtypic IAV infection is through rapid production of IL-10, which dampens innate inflammatory responses and limits lethal lung damage^22^. Notably, the vast majority of IL- 10+ memory T cells elicited by IAV infection also co-express IFN-γ^66^. To assess IFN-γ and IL-10 production by maternal memory T cells in offspring, we separately generated CD4.Cre x ROSA^tdTomato^ x IFN-γ^YFP^ (referred as T^Tom^ IFN-γ^YFP^) in which all conventional T cells express tdTomato and all IFN-γ expressing cells are YFP+; and CD4.Cre x ROSA^tdTomato^ x IL-10^GFP^ (T^Tom^ IL-10^GFP^) in which all conventional T cells express tdTomato and all IL-10 expressing cells are GFP+. Following X31 infection of reporter dams and PR8 challenge of Cre- (non T^Tom^) offspring, flow cytometric analysis of offspring lungs revealed that the majority of CD44^hi^ tdTom+ maternal-derived T cells were enriched for IFN-γ and IL-10 expression expression (**Fig S8C** and **S8D**).

In adult mice, memory T cell-derived IL-10 primarily acts on IL-10R-expressing innate immune cells to dampen lung inflammation and protect against secondary IAV infection^22^. We therefore asked whether maternally derived IAV-specific memory T cells confer protection to offspring through a similar mechanism. We first profiled IL-10R-expressing cells in the lungs of offspring born to X31-immune dams at day 2 following PR8 infection. IL-10R expression was predominantly detected among neutrophils and macrophages **(Fig S9A,B)**.

To test the functional importance of IL-10 signaling in offspring protection, we first blocked IL-10 receptor (IL-10R) signaling in offspring of X31_pre_ dams, which resulted in increased mortality in otherwise protected pups **(Fig 5E)**. To directly establish that IL-10 production by maternal T cells is necessary, we generated CD4.Cre × IL-10^f/f^ dams to specifically deplete IL-10 in T cells. Following X31 infection, dams with T cell-specific IL-10 deficiency similarly recovered as wild-type IL-10^f/f^ littermates dams **(Fig S9C)**, and were bred with wild-type sires at day 10 p.i. Offspring of these IL-10-deficient T cell dams trended toward increased susceptibility following PR8 infection **(Fig 5F)**, whereas offspring of X31-immune wild-type dams (IL-10^f/f^) were protected. Together, these findings suggest that maternal T cell-derived IL-10 mediates offspring protection against heterosubtypic infection.

Together, these findings support a model in which maternal memory T cells engage IL-10-responsive innate immune cells in the offspring lung and suppress pathogenic neutrophil-associated inflammation, thereby promoting disease tolerance during heterosubtypic IAV infection.

### Maternal vaccination protects offspring from lethal IAV infection in a T cell dependent manner

Natural infection and vaccination can elicit distinct T cell responses, resulting in potential differences in the magnitude and functional capacity of memory T cells during secondary challenge^67^. We therefore questioned if transgenerational protection by maternal memory T cells could be translationally leveraged through maternal vaccination using the licensed human FluMist vaccine. To do so, we investigated whether T cells generated following maternal vaccination with FluMist (2025–2026 formulation) could confer transgenerational protection from heterosubtypic infection.

Using HAI assays, we first validated that our 2025–2026 FluMist formulation does not elicit cross-reactive antibodies that neutralize PR8 similarly to previous formulations^19^ (**Fig 6A,B**). Consistent with these findings, intranasally vaccinated adult antibody-deficient μMT^-/-^ mice exhibited protection against lethal PR8 challenge, comparable to that observed in antibody-sufficient μMT^+/-^littermate controls (**Fig 6C, Fig S9A**). Together, these data establish that FluMist vaccination, similarly to X31 infection, can confer protection against secondary PR8 infection independently of antibody-mediated neutralization or other effector functions.

**Figure 6:**
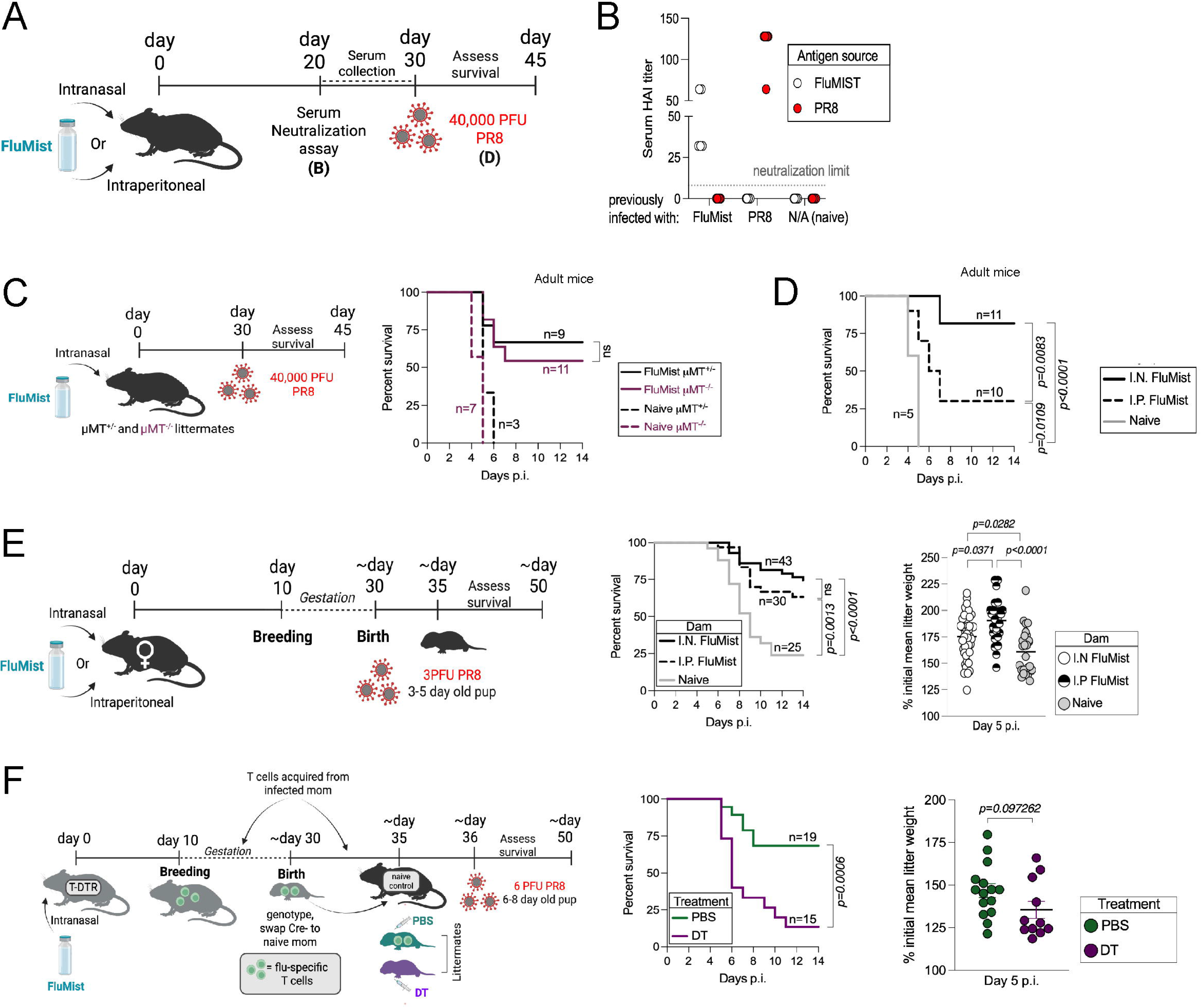
Maternal preconception FluMist vaccination confers transgenerational heterosubtypic protection to offspring in a T cell-dependent manner. **(A)** Experimental approach to validate the generation of protective T cell immunity in adult mice following FluMist vaccination. **(B)** Quantification of serum hemagglutination inhibition (HAI) titer of naïve, FluMist-vaccinated and PR8-infected adult mice (p.i.). The HAI was determined as the reciprocal of the last serum dilution to inhibit hemagglutination. N/A: non-infected; neutralization limit = 8. (C) Left panel: Experimental approach to test whether FluMist vaccination can protect B cell deficient mice. Right panel: Survival of naïve and FluMist-vaccinated μMT^+/-^ and μMT^-/-^ adult littermates following lethal PR8 infection. **(D)** Survival of littermate wild-type adult mice vaccinated with FluMist via the I.P or I.N routes. **(E)** Left panel: Experimental approach to test whether preconception maternal FluMist vaccination via intranasal (I.N) or intraperitoneal (I.P) route confers transgenerational heterosubtypic protection. Middle panel: Survival of offspring born to naïve, I.N or I.P vaccinated dams. Right panel: assessment of weight gain in the infected pups at day 5 p.i. **(F)** Left panel: Experimental approach to test whether maternal T cells generated following FluMist vaccination (represented in green) confer transgenerational protection. Middle panel: Survival of littermate offspring born from FluMist-immune T^DTR^ dams and given PBS (green) or DT (purple). T^DTR^: CD4.Cre^+/-^ x iDTR^lslFlox/wt^. Right panel: Percentage of initial body weight at d5 p.i. relative to the mean litter weight at d0. Each dot represents one mouse. Statistical significance for survival curves was determined by the log-rank test (Mantel-Cox). For weight comparisons, One-Way ANOVA (C-E), and Student’s t-test (G) were used.

Intranasal FluMist vaccination has been shown to promote superior heterosubtypic protection compared with systemic immunization, a difference attributed to the induction of lung-resident T_RM_. We confirmed that i.n. immunization favored the generation of IAV-specific CD8+ lung T_RM_, whereas intraperitoneal (i.p.) systemic immunization primarily generated circulating T_CM_/T_EM_ with minimal lung T_RM_ (**Fig S9B**). Additionally, i.n. immunized adult B6 exhibited significantly less morbidity and mortality following lethal PR8 challenge compared to i.p. immunized mice (**Fig 6D, Fig S9C**).

To understand whether maternal vaccination routes impact transgenerational immune defense, we immunized dams i.n or i.p. with FluMist, mated them with naïve sires, and challenged offspring with PR8 (**Fig 6E**, left). Offspring of vaccinated dams were significantly protected against lethal PR8 infection, regardless of maternal immunization route (**Fig 6E**, right). These results indicate that maternal lung T_RM_ are dispensable for transgenerational protection, consistent with our earlier finding that offspring protection is conferred irrespective of the tissue origin of transferred maternal T cells (**Fig 3I–J**). Finally, to confirm this vaccine-induced transgenerational protection is mediated by maternal T cells, we i.n. vaccinated T^DTR^ dams, mated them with naïve sires, and treated Cre-offspring with DT to deplete maternal T cells. DT treatment abrogated the protective benefit of maternal vaccination (**Fig 6F**), demonstrating that maternal T cells are required for offspring protection. Thus, vaccination strategies that elicit IAV-specific maternal T cells can protect offspring from lethal respiratory infection.

## Discussion

Maternal immune memory provides critical protection during early life, when susceptibility to severe infection is greatest. Although this protection has been largely attributed to maternally transferred antibodies, this form of defense is transient and readily circumvented by rapidly evolving pathogens such as influenza. Emerging studies in animal models suggest that maternal immune cells elicited by vaccination or infection can shape neonatal immune responses to pathogenic antigens^68,41,43,45,52,54^. Here, we reveal another facet of transgenerational immune memory transmission: the transplacental transfer of antigen-specific maternal memory T cells as a complementary mechanism of immune protection, enabling offspring to resist pathogens that escape maternal antibody-mediated defenses.

Our findings are consistent with earlier reports showing that maternal immunization with formalin-inactivated IAV during pregnancy confers heterosubtypic protection to offspring during the early life period ^69^. In addition, our cross-fostering studies indicate that protective maternal IAV-specific memory T cells for neonatal offspring are primarily acquired during gestation, in line with previous findings showing that Lm-OVA specific T cells transmitted to offspring *in utero*^52^. In contrast, using wild-type and IL-4R^-/-^ dams, previous studies reported that Th2 cells acquired from breastmilk of helminth-immune mothers can enhance immune responses of adult offspring against the same pathogen in an antigen-agnostic manner^45,54^. These results highlight both the differences in the mechanisms of T cell transfer as well as the timing of protection conferred to different pathogens, suggesting that transplacental and breastmilk-derived maternal T cells may have functionally different roles in offspring immunity. Future studies are required to decipher the molecular signals driving the transgenerational transmission of memory T cells as well as the contributions of different maternal T cell subsets to offspring protection.

Our work indicates that *in utero* transferred maternal memory T cells are particularly important during early life, when offspring T cell immunity is developmentally constrained. In both mice and humans, neonatal T cell responses exhibit innate-like features and are functionally distinct from those of adults^70^. Human infants fail to mount robust cytotoxic CD8+ T lymphocyte (CTL) responses required to efficiently clear respiratory syncytial virus (RSV) and influenza in the lower respiratory tract^71^. Mouse studies using PR8 and X31 showed that in neonates, the reduced CTL response stems from cell-intrinsic defects in TCR usage^5^, likely due to reduced expression of Terminal deoxynucleotidyl transferase (TdT), a specialized DNA polymerase that plays a critical role in generating TCR diversity^72^. However, important differences exist in immune ontogeny between mice and humans, particularly in the timing of peripheral T cell development. Fetal thymus-derived T cells populate peripheral tissues as early as the first trimester in humans, whereas this occurs only after birth in mice^73^. In our system, maternal memory T cells protected both neonatal and weanling offspring (the latter being more physiologically relevant to the human infant immune system), establishing that maternally acquired T cells can mediate defense both before and after establishment of the offspring peripheral T cell compartment. This interpretation is further supported by the preservation of protection in offspring lacking functional T cells, including TCRβ^-/-^ and TCR-restricted pups. Acquisition of maternal T cells may therefore provide an evolutionarily conserved benefit, encompassing critical immune components in immunodeficient offspring. This T cell-mediated compensatory protection is analogous to previously described maternal microchimeric myeloid cells providing complement factors in C1q^-/-^ pups to override *E. coli* infection^74^.

Our data indicate that maternal memory T cells protect offspring primarily by restraining early inflammatory dysregulation to limit lung damage. This mechanism is consistent with the well-established role of excessive innate immune activation in driving influenza-associated pathology. Elevated IL-1α/β drives early neutrophil recruitment and self-amplifying lung inflammation associated with poor outcomes in both mice and humans^75,76^. Likewise, increased MCP-1 correlates with disease severity in pediatric influenza cases^77^; whereas type I interferon overproduction is a hallmark of neonatal influenza susceptibility^78^. Excessive IL-27 production has also been implicated in promoting immunopathology during the early stages of influenza infection in adults^79^. Notably, multiple studies have described an association between maternal microchimerism at birth and protection from hospitalization and severe disease in human infants, including respiratory tract infection, non-malarial fever, and symptomatic malaria^46,80,81^. Together, these observations support a model in which maternally transferred memory T cells enhance disease tolerance by preventing the early inflammatory cascade that underlies severe (and potentially lethal) infection-induced disease.

The ability of maternal memory T cells to confer heterosubtypic protection suggests that they can protect offspring not only against the maternal infecting strain but also against related pathogens sharing conserved antigens. This broad reactivity mirrors observations in humans, where pre-existing CD4 and CD8 T cells recognizing conserved influenza epitopes correlate with reduced viral shedding and disease severity following infection with antigenically distinct strains, even in the absence of cross-neutralizing antibodies^82,83^. Such breadth has important implications for maternal vaccination, particularly because infants are among the populations most susceptible to severe respiratory infections and cannot receive live attenuated influenza vaccination before six months of age^36^. IAV vaccination during pregnancy substantially reduces infant hospitalization, especially during the first six months of life^84,85^, and maternal vaccination with the RSV prefusion F antigen (RSVpreF) similarly lowers RSV-associated hospitalization in early infancy^86,87^. These benefits have been largely attributed to maternal antibodies; however, the contribution of maternally transferred immune cells has remained largely unexplored. Notably, together with the enrichment of SARS-CoV-2 Spike-specific T cells in breastmilk following maternal mRNA vaccination maternal^14^, our findings support the concept that maternal memory T cells may represent an unrecognized component of vaccine-induced immunity transmitted across generations.

Our study expands the current paradigm of maternal immune protection by identifying antigen-specific maternal memory T cells as a previously unrecognized mechanism of transgenerational immunity against respiratory infection. By complementing the protective capacity of maternal antibodies, maternally transferred memory T cells provide broad protection against pathogens that evade humoral immunity, thereby extending the spectrum of immune defenses available to the offspring during a period of heightened vulnerability, as well as providing a unique function in protection against immunopathology. These findings refine the maternal–offspring immune partnership and establish transferred maternal memory T cells as an evolutionarily conserved layer of immune protection with important implications for the design of next-generation maternal vaccines.

## Limitations of the Study

One limitation of this study is that murine and human immune ontogeny differ substantially, particularly with respect to the timing of T cell development and maturation. To partially address this, we evaluated offspring protection in both neonatal and weanling mice, the latter more closely approximating the developmental stage of the human infant immune system. Nevertheless, whether maternal antigen-specific T cells confer comparable protection in humans remains to be determined. Further, inherited maternal T cells are rare, making detailed phenotypic characterization challenging. However, even at low frequency, these cells appear to exert a profound influence on offspring immunity and susceptibility.

## Acknowledgments

This work was supported by the Pew Charitable Trusts (M.A.K.), the NIH (R01 R01A1173199 to M.A.K.). H.T. is a Washington Research Foundation (WRF) Postdoctoral Fellow. BNJ is supported by a Cell Molecular Biology Training Grant (T32 GM136534) from the University of Washington. M.E.C is supported by a F31 fellowship (1F31AI200407) from the NIH. Additional support was provided by NIH P30 CA015704 of the Fred Hutch/University of Washington/Seattle Children’s Cancer Consortium, which includes the Genomics & Bioinformatics Shared Resource (RRID:SCR_022606), Comparative Medicine Shared Resource (RRID:SCR_022610), Experimental Histopathology Core Facility (RRID:SCR_022612), and Flow Cytometry Core Facility (RRID:SCR_022613). X31 and PR8 strains were kindly provided by Dr. Paul Thomas (Fred Hutch). CD8 tetramers (IAV NP366-372, and SIINFEKL peptide corresponding to OVA257-264) were obtained from the NIH tetramer core (NTCF). IAV NP311-325 CD4 tetramer was kindly provided by Dr Thomas Griffith^88^. AstraZeneca provided FluMist through a material transfer agreement. The authors would also like to acknowledge the scientific services staff at Fred Hutch. The content is solely the responsibility of the authors and does not necessarily represent the official views of the National Institutes of Health.

## Author contributions

H.T., W.E.H, and M.A.K. conceived and designed the experiments.

The experiments were performed by H.T with help from B.N.J, M.E.C, E.E.L, D.M.R. and S.A.

H.T. and M.A.K. prepared the early manuscript drafts, and all authors contributed thereafter.

## Declaration of generative AI and AI-assisted technologies in the writing process

During the preparation of this work, the authors used Claude and ChatGPT in order to proof-read the manuscript for grammatical errors and to improve the overall readability. After using these tools, the authors reviewed and edited the content as needed and take full responsibility for the content of the published article.

## Materials and Methods

### Mice

Specific pathogen-free (SPF) C57Bl/6J (Jax000664), Thy1.1 (*B6.PL-Thy1a/CyJ*, Jax000406), CD45.1 (*B6.SJL-Ptprc^a^ Pepc^b^/BoyJ*, Jax002014), mMT^-/-^ (B6.129S2*-Ighm^tm1Cgn^/J,* Jax002288), TCRbd^-/-^, (B6.129P2-*Tcrb^tm1Mom^ Tcrd^tm1Mom^*/J, Jax002122), OT-1 (*C57BL/6- Tg(TcraTcrb)1100Mjb/J*, Jax003831), OT-2 (*B6.Cg-Tg(TcraTcrb)425Cbn/J*, Jax004194), CD4.Cre (*B6.Cg-Tg(Cd4-cre)1Cwi/BfluJ*, Jax022071), CD4.CreER^T2^ (*B6(129X1)-Tg(Cd4-cre/ERT2)11Gnri/J*, Jax 022356), iDTR^flox/flox^ (*C57BL/6-Gt(ROSA)26Sor^tm1(HBEGF)Awai^/J*, Jax007900), IL-10^flox/flox^ (*B6.129P2(C)-Il10^tm1Roer^/MbogJ*, Jax036598), ZsGreen^flox/flox^ (*B6.Cg- Gt(ROSA)26Sor^tm6(CAG-ZsGreen1)Hze^/J*, Jax007906), and tdTomato^flox/flox^ (*B6.Cg- Gt(ROSA)26Sor^tm14(CAG-tdTomato)Hze^/J*, Jax007914) mice were purchased from the Jackson Laboratory. SPF C57BL/6NTac (B6-F) and BALB/c (BALB/cAnNTac) mice were purchased from Taconic Biosciences. IFNg^YFP^ (GREAT, *B6.129S4-Ifng^tm3.1Lky^/J*, Jax017581) mice were provided by Dr. Jarrod Dudakov (Fred Hutch Cancer Center), and IL10^GFP^ (Tiger, *B6.129S6-Il10^tm1Flv^/J*, Jax008379) mice were obtained from Dr Geoffrey Hill (Fred Hutch Cancer Center). All mice were bred and maintained at an American Association for the Accreditation of Laboratory Animal Care (AALAC)-accredited animal facility at the Fred Hutchinson Cancer Center (RRID:SCR_022610) under SPF conditions, and housed in accordance with the procedures outlined in the *Guide for the Care and Use of Laboratory Animals*. All experiments with mice were performed in accordance with the guidelines of the Animal Care and Use Committee at the Fred Hutchinson Cancer Center. All mice were maintained under a 12-h light-dark cycle (7 a.m. to 7 p.m.) and given a standard chow diet (Teklad 2918). All mice were kept on neutral water (pH 6.5-7.5) *ad libitum* except for the TCRbd^-/-^mice that were on acidic pH water (pH 2.5-3.5).

Dams of the same experimental conditions were bred for 3-4 nights and were then regrouped together for approximately 15 days (until late pregnancy) to homogenate their microbiota. Pregnant females were separated and housed individually until parturition. Age matched offspring of both sexes were used in experiments. Only nulliparous dams were used for any experiment involving offspring survival, unless specified. In some cases, pups were tattooed at birth via 25G needle (Ketchum).

### Influenza infection

The mouse-adapted PR8 (H1N1) and X31 (H3N2) strains were generously obtained from Dr. Paul Thomas (via Dr. Andrew Oberst, UW). Individual aliquots from the same batch of viruses, cultured in fertilized chicken egg, were kept at -80C until use. Aliquots were thawed at room temperature for 5 minutes and serially diluted in sterile PBS to get the suitable concentrations and kept on ice until administration in mice in an accredited Biosafety level 2 (BSL2) mouse husbandry room. Mice were anesthetized with isoflurane and intranasally infected with Influenza. For sublethal infections, adult mice were infected with 1,000 plaque forming units (PFU) X31 or 10 PFU PR8 diluted in 30µl PBS. For PR8 lethal challenges, adult mice were intranasally infected with 5,000 PFU or 40,000 PFU in 30µl PBS. Weights were daily recorded following infection, and mice losing more than 20% of initial body weight were humanely euthanized, as approved by the Fred Hutch Institutional Animal Care and Use Committee and Institutional Review Board. Male and female weanlings (18-21 days old) were separated from dams, supplemented with dietary recovery gel (ClearH_2_O 72-06-5022), and infected with 150 PFU PR8 in 10µl PBS on the same day. Neonatal pups (3-5 days old) were infected with 3 PFU PR8 (B6) or 12 PFU PR8 (F1) in 3µl PBS.

### Listeria monocytogenes culture, infection and bacterial load quantification

The 10403s strain of *Listeria monocytogenes* (naturally resistant to streptomycin) expressing chromosomally integrated ovalbumin (Lm-OVA) was obtained from Dr Olivier Harrison. Lm-OVA was grown at 37°C in Brain Heart Infusion (BHI – Sigma-Aldrich 53286) broth containing 200 µg/mL streptomycin under aerobic condition until reaching exponential growth phase (OD_600_ 0.6-0.8). Bacterial cultures were washed three times with PBS, and centrifuged to obtain concentrated aliquots, which were frozen at -80°C in 10% glycerol until use. One day after storage, an aliquot of Lm-OVA was thawed on ice, 10-fold serially diluted in water, plated on BHI agar (200 µg/mL streptomycin), and incubated at 37°C overnight to determine bacterial concentration of the batch. For Lm-OVA infection, a vial of the batch was thawed on ice and serially diluted to obtain the desired concentration and kept on ice until administration in mice on the same day. Adult mice were anesthetized with isoflurane and infected with 30µl Lm-OVA in PBS. For experiments involving offspring survival, dams were infected with 2x10^6^ CFU Lm-OVA. To determine *in vivo* bacterial load at day 10 post-infection, lungs, liver, and feces were collected following euthanasia by CO_2_ and homogenized at 100 mg/mL in PBS, serially diluted, plated on BHI agar (200 µg/mL streptomycin), and incubated at 37°C overnight.

### Immunization with FLuMIST and PoIyI:C+OVA

Live attenuated Influenza vaccine, FluMist (2025-2026 formulation) was obtained from AstraZeneca (NDC 66019-112-00). Mice were immunized either intraperitoneally or intranasally with 10^5.5-6.5^ fluorescent focus units (FFU), corresponding to 1/10^th^ of the human dose ^19^. For PoIyI:C+OVA immunization, mice were anesthetized with isoflurane and received intranasal administration of 50µl PBS containing 100µg PolyI:C LMW (Invitrogen NC0131768 TLRL-PICW-250) and 3.33 mg grade III OVA (Sigma-Aldrich A5378-25G).

### Milk and serum collection

Lactating dams were separated from pups for 3 hours at day 3 post-parturition and anesthetized via nose-cone with isoflurane. Dams were injected intraperitoneally with 250µL of PBS containing 2 IU oxytocin (Alfa Aesar J63421), and milk production was stimulated with hand massage. Milk was collected using microhematocrit capillary tubes, transferred to a microfuge tube. Milk was then diluted 1/3 in PBS, centrifuged 3 times to clear the aqueous phase that was stored at -80°C for subsequent analysis. For serum collection from adult mice, blood was collected via saphenous vein bleed or cardiac puncture of mice at the time of euthanasia. Blood from neonates was collected after decapitation. Blood was clotted at room temperature for 30-60min, centrifuged at 4°C, 13,000*g* for 30min, and the serum supernatant was transferred to a separate microfuge tube and kept at -80°C for subsequent analysis.

### Cytokine quantification

Cytokines were quantified in breastmilk, serum, and lungs. Lungs were homogenized at 100 mg/mL in PBS containing protease inhibitor (Roche 4693116001), centrifuged at 13,000g for 10min and supernatant was used. Cytokines were quantified using the LEGENDplex Mouse Inflammation Panel (BioLegend 740150) following the manufacturer’s instructions.

### Hemagglutination inhibition assay (HAI)

The hemagglutinin (HA) titer of our FluMist, X31 and PR8 stocks were determined by incubating 25 µl of different 2-fold dilutions of viral stock with 25 µl of 0.5% chicken red blood cells (cRBC) (Lampire, Fisher Scientific 50-414-455) for 1 hour at room temperature. The HA titer was defined as the highest viral stock dilution that caused hemagglutination with cRBC. To determine serum HAI titers, serum from naïve, PR8, X31, or FluMist-immune mice were diluted at ¼ (v/v) in Receptor Destroying Enzyme II (RDE) (Accurate Chemical YCC340122) and incubated 18-20hours at 37°C prior to HAI assay to eliminate non-specific serum inhibitors. The reaction was stopped by incubating the serum/RDE II mixture at 56°C for 1h to destroy the RDE II enzyme. HAI assay was performed at room temperature by incubating 25 µl of 2-fold serial dilutions of treated serum with 25 µl of 4HA units for 15 minutes, then incubating this mixture with 50 µl of 0.5% cRBC for 1 hour. The HAI titer was determined as the reciprocal of the highest serum dilution that prevented hemagglutination of viral antigens with cRBC.

### Serum adoptive transfer

For serum transfer in neonates (p3-p5), sera were collected from nulliparous or parous X31-immune or PR8-immune dams and filtered through a 0.22 µm filter (Millex, cat# SLGV013SL) on the day of transfer. Pups were injected intragastrically with 30-40 µl of filtered serum or PBS using Hamilton gas syringe (HAM7656-01-1EA) and 30G needle, and infected with 30 PFU X31 or 3PFU PR8 on the same day. For adults, naïve dams were retroorbitally injected with 100 µl of filtered serum freshly collected from X31-immune dams (day 10 p.i.) at the same day of transfer. The recipient dams were mated with B6 sires on the same day to produce progeny that was subsequently infected for survival purpose.

### Diphtheria toxin administration

Adult mice were intraperitoneally injected with Diphtheria toxin (DT) (Sigma D0564) on two consecutive days. They were treated with 30ng/g of body weight on the first day and 10ng/g on the second day. Neonatal pups (p5-8) received a single intragastric injection of 50ng/g of body weight.

### Antibody treatments

Pups received single intragastric injections of antibodies at day 1 p.i. with PR8. For IL-10R blockade, pups received 150µg of anti-IL-10R (BioXCell, BE0050, clone 1B1.3A) or 150µg of rat IgG1 isotype control (BioXCell, BE0088, clone HRPN). Neutrophils were depleted using a dual antibody injection strategy ^89,90^, by combining 380µg of anti-Ly6G (BioXCell, BE0075-1, clone 1A8) with 160µg of anti-rat Igκ (BioXCell, BE0122, clone MAR18.5). The control littermates received 380µg of rat IgG2a isotype (BioXCell, BP0085, clone C1.18.4).

### Viral load quantification

Total RNA was extracted from the left lung lobe at the indicated timepoints. Lungs were collected and homogenized in TRIzol (Thermo 15596018) and RNA was enriched by phase separation using chloroform (1/5 -v/v) followed by centrifugation at 12,000 × g at 4°C for 10 minutes. The RNA was enriched either by Isopropanol precipitation followed by 3X wash with 70% Ethanol or using Zymo RNA Miniprep kit (Zymo R2050). The purified RNA was treated with Turbo DNA-free kit (Invitrogen AM1907) to remove genomic DNA. The treated RNA was reverse transcripted into cDNA using iScript™ cDNA Synthesis Kit (BioRad 1708890). IAV viral load was quantified by qPCR targeting *NS1* transcripts in a total reaction volume of 20µl using Bio-rad iTaq Universal SYBR Green Supermix (172-5122). The reaction contained 10µl SYBR Green + 1µl cDNA of (20ng) + 1µl l Forward primer (10µM; TGTCAAGCTTTCAGGTAGATTG) + 1µl Reverse primer (10µM; CTCTTAGGGATTTCTGATCTC) +7µl DNase/RNase free water. The β-actin transcript was used for normalization (Forward: CACTGTCGAGTCGCGTCC; Reverse: TCATCCATGGCGAACTGGTG). The amplification cycle was: 1X (95°C C 3min) followed by 40X (95°C 15sec, 59°C 40sec, Plate read) in a Biorad CFX96 qPCR machine. The ΔΔCT method was used for relative quantification.

### Maternal T cell detection in offspring lungs by Countable PCR

Presence of maternal-inherited zsGreen-expressing T cells was detected in the lungs by Countable PCR targeting *zsGreen1* transcripts. The reaction was performed in a total volume of 50µl using Countable Labs 4X Countable PCR mix (Countable Labs KT0004) and Universal Multiplex probes (Countable Labs KT0005). The reaction contained 12.5µl 4X PCR mix + 35µl cDNA (800ng/uL) + 1µl 50X UM Primer mix (10µM; Forward, with UM sequence underlined: <u>CAGAAGACGGCATACGAGAT</u>AACCAGAAGTGGCACCTGAC; Reverse: GGCATTAAAGCAGCGTATCC) + 1µl 50X UM-02 Probe + 0.5µl DNase/RNase free water. The amplification cycle was: 1X (95°C C 20sec) followed by 10 touchdown cycles (65°C to 60°C), then 39X (95°C 15sec, 60°C 1min). zsGreen-specific counts were imaged and quantified by the Countable PCR system.

### Nanostring gene expression quantification

Nanostring assays were performed by the Fred Hutch Genomics & Bioinformatics Shared Resource (RRID:SCR_022606) on samples with a RIN>8. 100 ng of purified total RNA in 5µL was utilized as input into NanoString’s nCounter Mouse Immunology V1 panel (531-115000052). The CodeSet reference was GX (CP9231X2). Samples were processed according to the manufacturer’s protocol. RNA was hybridized with the mouse immunology code-set for 16 hours at 65°C. Target-probe complexes were processed and immobilized to an nCounter cartridge on the nCounter Prep Station and then transferred to the nCounter MAX/FLEX system to quantify probe abundances. Sample QC and reporter probe counts were then analyzed using the nSolver Analysis Software 4.0. The mRNA panel, nCounter system, and nSolver software were manufactured by Nanostring Technologies, Inc., Seattle, WA.

### Total RNA sequencing

For RNA-Seq, only samples with a RIN>8 were included. RNA-Seq was performed by Plasmidsaurus using Illumina Sequencing Technology using Illumina NovaSeq X Plus (catalog no. 20084804) and a 3’ end counting approach with custom analysis and annotation. For sequencing library preparation, mRNA was converted into complementary cDNA via reverse transcription and second-strand synthesis, followed by tagmentation, library indexing, and amplification. The following pipeline was applied for bioinformatics analysis: (1) FastQ generation and demux with BCL Convert v4.3.6 and fqtk v0.3.1.; (2) Read-filtering using FastP v0.24.0: poly-X tail trimming, 3’ quality-based tail trimming, a minimum Phred quality score of 15, and a minimum length requirement of 50 bp.; (3) Alignment to the appropriate reference genome using STAR aligner v2.7.* with non-canonical splice junction removal and output of unmapped reads.; (4) Coordinate sorting of BAM files using samtools v1.21.; (5) UMI based de-duplication: Removal of PCR and optical duplicates using UMICollapse v1.1.0.; (6) Mapping QC: Alignment quality metrics, strand specificity, and read distribution across genomic features using RustQC v0.2.1.; (7) Generation of comprehensive QC report using MultiQC v1.33.; (8) Gene-expression quantification using featureCounts (subread package v2.1.1) with strand-specific counting, multi-mapping read fractional assignment, exons and three prime UTR as the feature identifiers, and grouped by gene_id. Final gene counts were annotated with gene biotype and other metadata extracted from the reference GTF file.; (9) Sample-sample correlations for sample-sample heatmap and PCA were calculated on normalized counts (TMM, trimmed mean of M-values) using Pearson correlation.; (10) Differential expression using edgePython v0.2.5.; (11) Functional enrichment performed for human and mouse samples using gene set enrichment analysis with GSEApy v0.12 using the MSigDB Hallmark gene set.

### Histopathology

Pups were euthanized by 2.5% avertin overdose and lungs were inflated with 10% neutral-buffered formalin solution (NBF). Individual inflated lungs were fixed in a pre-filled container with 10% NBF (Simport M961) and submitted to Fred Hutch Histopathology core (RRID:SCR_022612). Lungs sections were stained with hematoxylin and eosin (H&E) and were scored on a scale of 0-4 based on 3 parameters: percentage of lung affected, airway inflammation and interstitial inflammation. Other features included necrotic debris, bronchiolar epithelial necrosis, fibrin, alveolar proteinosis, and BALT hyperplasia. Scoring was performed with groups blinded to a board-certified pathologist and groups were unblinded after scoring.

### Tissue processing

Neonatal mice (6 to 10 days old) were euthanized by intragastric injection of 100µl of 2.5% avertin followed by decapitation prior to tissue collection. Adult mice were euthanized by intraperitoneal injection of 1ml 2.5% avertin overdose followed by cervical dislocation. In some cases, neonates received retroorbital injection of 0.75µg anti-CD45-BUV395 (Fischer Scientific 363-0451-82, clone 30-F11) and adults received retroorbital injection of 3µg of anti-CD45-BUV395, anti-CD45.2-APC (Tonbo SKU 20-0454-U100, clone 104) or anti-CD45-APC (Tonbo SKU 20-0451-U100, clone 30-F11) 5 minutes after avertin injection to label leukocytes in the vasculature ^64^. Adult lungs were place in cold DMEM (Gibco) and neonatal lungs were collected in cold DMEM (Gibco) supplemented with with 2mM L-glutamine, 1mM sodium pyruvate, 100U/ml penicillin, 100mg/mL streptomycin (all from Gibco) and 3% (v/v) Fetal Calf Serum (FCS; Cytiva) in in C-Tubes (Miltenyi). Spleens were placed in cold complete media (Gibco RPMI 1640 supplemented with 2mM L-glutamine, 1mM sodium pyruvate, 100U/ml penicillin, 100mg/mL streptomycin) supplemented with 3% (v/v) FCS in in C-Tubes. Lungs were briefly dissociated with a GentleMACS Dissociator (Miltenyi) before to be incubated at 37 °C for 30 min in 5mL collagenase-Type IV (10 µg/ml, Sigma-Aldrich) and 10 µg/ml DNaseI (Roche) for enzymatic dissociation. Spleens were incubated in 5mL complete media (without FCS) containing 5µg/ml DNaseI and 1mg/mL Collagenase IV for 30min at 37°C. Tissues were homogenized GentleMACS Dissociator (Miltenyi) to get single cell suspensions. The enzymatic reaction was stopped with the addition of 10 ml cold complete media and tissue homogenate was passed through a 70μm Filter (Fisher). Cells were pelleted at 475x*g* and resuspended in 1 ml ACK lysing buffer (Gibco A10492-01) for 10 min at RT to lyse the red blood cells and neutralized using cold complete media. Cells were next pelleted at 475x*g* and the pellet was directly resuspended in complete media, EasySep buffer (StemCell 19856) or PBS containing 0.5% BSA and 2mM EDTA for downstream analysis or T cell isolation.

### T cell isolation and adoptive transfer

T cells were isolated from single cell suspensions of the lungs or spleen. For experiments involving adoptive transfers, T cells were isolated using Miltenyi positive selection beads for CD8+ T cells (130-117-044), and CD4+ T cells (130-117-043) following the manufacturer’s instructions. 5 million CD4+ and 5 million CD8+ T cells from the spleen in 200μl PBS were retroorbitally transferred in naïve recipient dams; 0.5 million CD4+ and 1 million CD8+ T cells from the lungs in 200μl PBS were retroorbitally transferred in naïve recipient dams. The recipient dams were mated with B6 sires on the same day to produce progeny that was subsequently infected for survival purpose. To track maternal-acquired fluorescent T cells in neonates by flow cytometry, total T cells were isolated either using Miltenyi CD4+ and CD8+ positive selection beads or negative selection kit from StemCell (19851) following the manufacturer’s instructions prior to staining for flow cytometric analysis.

### Tetramer staining

Tetramers were received from the NIH tetramer core facility (NTCF). Fluorescently conjugated MHC-I coupled to ASNENMETM peptide corresponding to IAV NP_366-374_ and MHC-I coupled to SIINFEKL peptide corresponding to OVA_257-264_ were used to track Influenza and OVA-specific CD8 T cells respectively. Fluorescently conjugated mouse CD1d loaded with PBS-57 was used to identify iNKT cells ^91^. APC-conjugated, MHC-II coupled QVYSLIRPNENPAHK (NP_311-325_) CD4 tetramer was received from Dr Marion Pepper (UW). Whole lungs or spleen single cell suspensions or T cells isolated using the StemCell negative selection kit were stained with 5.5 µg/ml NP_366-374_ or 10 µg/ml NP_311-325_ at room temperature for 1 hour or 2.5 µg/ml OVA_257-264_ at 37C for 30 minutes. Cells were then washed and stained with surface marker antibodies for flow cytometry. In some cases, to enrich for tetramer specific T cells, cells were stained with NP_366-374_ and NP_366-374_ tetramers conjugated to APC or PE and before to be washed and incubated with 25 μl anti-APC (Miltenyi Biotec 130-090-855) and/or anti-PE (Miltenyi Biotec 130-105-639) microbeads at 4C for 30 min. Tetramer-positive cells were enriched by magnetic selection using LS columns (Miltenyi Biotec 130-042-401) as previously described ^92,93^. The enriched fraction and the flow-through (non-enriched fraction) were then separately stained with surface marker antibodies for flow cytometic analysis.

### Universal LIPSTIC labeling

At 2 days post-infection with 3PFU PR8, neonatal mice (6-7 days old) were injected systemically twice (1 hour apart) with 30µl of 100mM Biotin-Ahx-LPETGS-NH2 substrate (LifeTein 5467). The lungs were collected 1 hour after the second injection, after euthanasia using 2.5% avertin overdose followed by decapitation, and the tissues were dissociated to prepare single cell suspensions for flow cytometry analysis. The cells at proximity of the donor cells were determined as biotin+ following staining with anti-biotin APC (Miltenyi 130-113-288, final dilution 1/50).

### Flow cytometry

2-5x10^6^ cells were incubated for 15 minutes at room temperature in 55 μl of fixable Ghost Dye Violet 510 (Tonbo 13-0870-T100) or Ghost Dye Red 780 (Tonbo 13-0865-T500) and Fc Shield (1/200 – Tonbo 70-0161-U100) diluted in FACS buffer (PBS supplemented with 2% (v/v) FCS and 2mM EDTA; Sigma-Aldrich). 55 μl of the extracellular antibody stain diluted in FACS buffer was added and cells were incubated for 30min at 4°C. After staining, 100μl FACS buffer was added, and cells were centrifuged at 475*g* for 5min. For intracellular staining, cells were fixed and permeabilized using the Foxp3/Transcription Factor Staining Buffer Set (ThermoFisher Scientific 00-5523-00) for 30min at room temperature. Cells were centrifuged at 685*g* for 5min and washed with 200μl of Foxp3 Perm/Wash buffer and centrifuged at 685*g* for 5min. The pellet was then resuspended in intracellular antibody staining mix and incubated at room temperature for 1hr. Cells were washed with 150μl Perm/Wash, centrifuged at 685*g* for 5min, and resuspended in 200 μl of Perm/Wash. The stained cells were filtered with a FACS plate or tube strainer, and Accucheck Counting Beads (Invitrogen) were added on each sample prior to FACS analysis using a BD FACSymphony A5 High-Parameter Cell Analyzer or a Cytek Aurora spectral cytometer (Fred Hutch Cytometry Core, RRID:SCR_022613). Beads (UltraComp eBeads or Spectral UltraComp eBeads, Invitrogen) stained with individual antibodies were used for compensation. zsGreen1+, tdTOM+, GFP+, or YFP+ blood lymphocytes were used for compensation. The list of antibody clones/dyes and concentrations used are detailed in the Key resources Table. All data were analyzed in FlowJo, version 10.

### Statistics

Statistical significance was determined as indicated in the figure legends with Prism 10 (GraphPad Software Inc).

## Key resources table

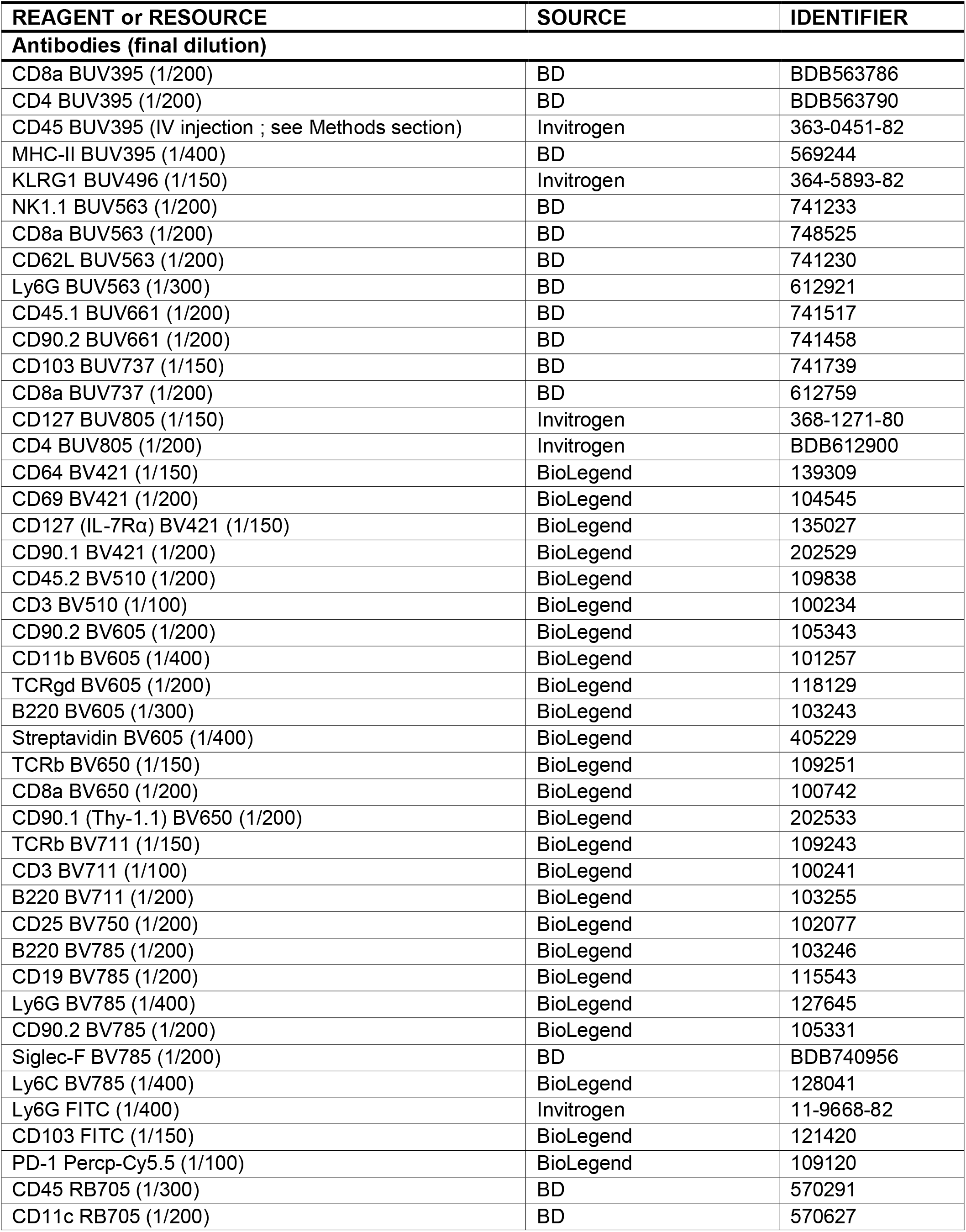
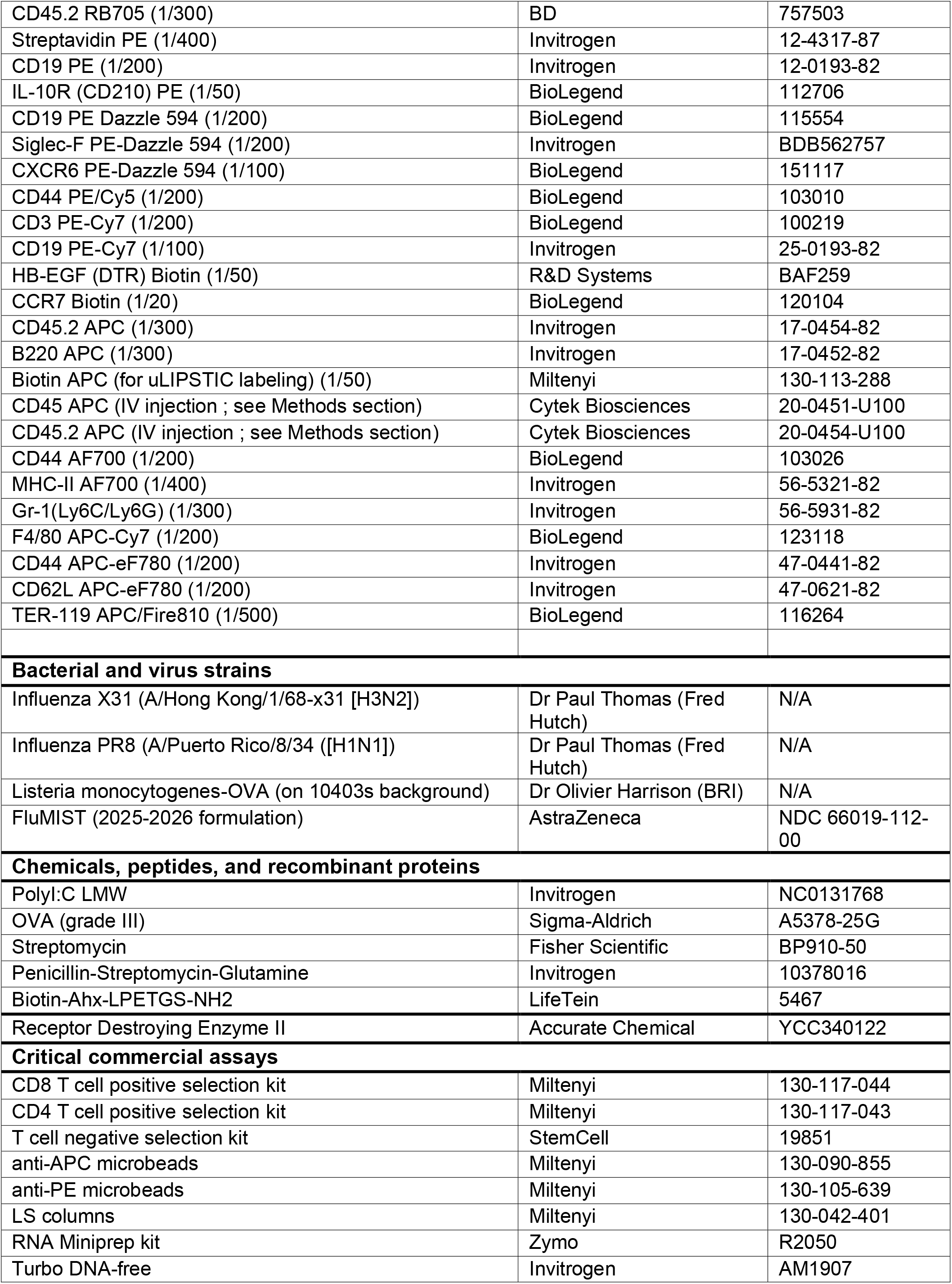
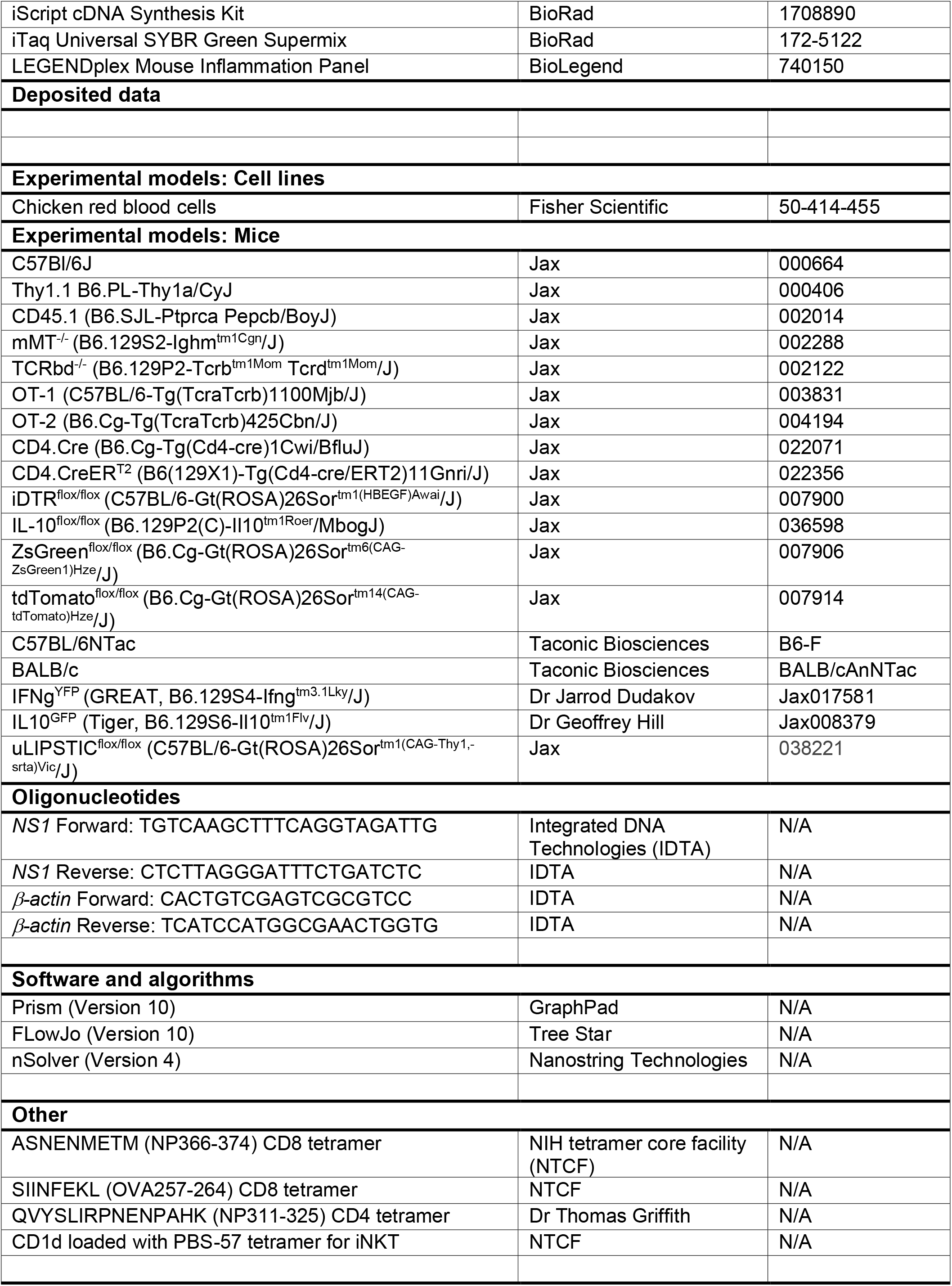

## Supplemental Figure

**Figure S1: Maternal preconception infection with IAV leads maternal IAV-specific T cell immunity and offspring protection against heterosubtypic infection. (A)** Weight loss assessment in female C57BL/6J mice following infection with 1,000 PFU X31 or 10 PFU PR8. **(B)** Relative X31 (left panel) or PR8 (viral load) in the lungs at day 1, 5, and 10 post-infection (p.i.). Viral *NS1* transcripts levels were normalized to β-*actin*. **(C)** Flow cytometric analysis of T cells responses of naïve and X31 or PR8-infected dams at day 10 p.i.; IV: anti-CD45 antibody injected intravenously 5 minutes prior to tissue collection. **(D)** Experimental approach (left panel) and weight gain of pups born to naïve or X31-immune (X31_pre_) C57BL/6NTac dams and infected with PR8 at days 3 and 5 p.i. **(E)** Experimental approach (left panel), survival (middle panel), and weight gain of pups born to naïve or X31_pre_ C57BL/6J dams and infected with PR8 at days 3 and 5 p.i. **(F)** Survival of F1 hybrid pups born to naïve C57BL/6NTac dams crossed to BALB/c sires. Pups were infected with 3, 9, 12, 15 or 30 PFU PR8 between day 3-5 post birth. Each dot represents an individual pup in B-E. Statistical significance for survival curves was determined by the log-rank test (Mantel-Cox), and non-parametric Student’s t-test (Mann-Whitney) for weight gain comparisons in D and E.

**Figure S2: Maternal preconception infection with IAV protects weanling but not adult offspring against heterosubtypic infection. (A)** Experimental approach and weight gain assessment of weanling males and females born to naïve or X31_pre_ C57BL/6J dams and infected with 150 PFU PR8 between day 18-20 post birth. **(B)** Experimental approach and weight gain assessment of adult males and females pups born to naïve or X31_pre_ C57BL/6J dams and infected with 5,000 PFU PR8 between weeks 7-9 post birth. Statistical significance for survival curves was determined by the log-rank test (Mantel-Cox), and Student’s t-test for morbidity comparisons.

**Figure S3: Maternal serum soluble factors are dispensable for transgenerational IAV heterosubtypic protection. (A)** Cytokine levels (day 3 post-partum) in the serum of naïve and X31_pre_ dams. The magenta boxes represent cytokines with heightened levels in X31_pre_ dams. **(B)** Experimental approach (left panel) showing serum collection from PR8-immune mice at day 30 p.i. and serum adoptive transfer in naïve recipient pups on the same day. p3-5 recipients were infected with PR8 (right panel) and survival was monitored. Statistical significance for survival curves was determined by the log-rank test (Mantel-Cox), and non-parametric Student’s t-test (Mann-Whitney) for cytokine level comparisons.

**Figure S4: Validation of the utility of the T^DTR-zsGreen^ model to efficiently deplete maternal T cells in the offspring. (A)** Flow cytometric confirmation of DTR expression by T cells in the blood of T^DTR^ mice. T^DTR^: CD4.Cre^+/-^x iDTR^lslFlox/wt^. **(B)** Experimental approach (left) and Flow cytometric confirmation (right) of zsGreen+ T cells depletion in the blood of T^DTR-zsGreen^ mice following Diphtheria toxin (DT) administration. T^DTR-zsGreen^: CD4.Cre^+/-^x iDTR^lslFlox/wt^ x zsGreen^lslFlox/wt^. **(C)** Experimental approach (left) and Flow cytometric confirmation (right) of the depletion of adoptively transferred CD45.2+ CD90.2+ zsGreen+ T cells splenocytes from T^DTR-zsGreen^ mice. The spleen of CD45.1+ CD90.1+ recipient pups were analyzed on the day following DT or PBS injection. **(D)** Analysis of donor T cells, B cells, and recipient T cell frequencies out of total live recipient splenocytes on the day following DT or PBS injection. Statistical significance was determined by Student’s t-test.

**Figure S5: Maternal IAV-specific T cells are sufficient to confer heterosubtypic protection in offspring. (A)** Experimental approach and flow cytometric analysis of T cell responses in the lungs and the spleen of adult mice following intranasal immunization with Poly(I:C)+OVA or infection with X31 or Lm-OVA at day 10 p.i.. **(B)** Representative flow plots of the lungs T cell responses of Poly(I:C)+OVA (upper panel) and Poly(I:C)+X31 (bottom panel) immune dams at day 10 p.i. **(C)** Experimental approach (left) and assessment of weight loss in female mice infected with Lm-OVA (middle) and bacterial clearance at day 10 p.i. in the lungs, liver, and feces. **(D)** Experimental approach to test whether preconception maternal infection with Lm-OVA protects offspring against PR8 infection. **(E)** Representative flow plots of the lungs T cell responses of Lm-OVA (upper panel) and X31 (bottom panel) immune dams at day 10 p.i. **(F)** Survival of pups born to naïve, Lm-OVA or X31-immune dams and infected with PR8 between day 3-5 post birth. **(G)** Representative flow plots of isolated T cells from the lungs and the spleen of X31 (left panel) and Poly(I:C)+OVA (right panel) immune dams at day 10 p.i. Pre-gate: CD45+ > Ter119-B220-CD19-Siglec-F-Ly6G-> CD90.2+ TCRβ+. These T cells were adoptively transferred in naïve recipient dams. Statistical significance for survival curves was determined by the log-rank test (Mantel-Cox), and by One-Way ANOVA for weight gain comparisons in F.

**Figure S6: Maternal memory T cells confer transgenerational heterosubtypic protection by limiting inflammatory responses during early timing of infection. (A)** Left panel: Representative lungs H&E staining at day 5 post PR8 infection of pups born from naïve (right) or X31_pre_ dams (left). Middle panel: Cumulative lungs pathology score. Right panel: Percentage of lungs area affected by the pathology; n=10 mice per group. **(B-D)** Cytokine levels in the lungs and serum of pups born from naïve or X31_pre_ dams at steady state (D), or at day 3 (B) or day 5 post-PR8 infection (C). **(E)** Volcano plot of differential expression of immune genes in the lungs of uninfected offspring of naïve or X31_pre_ dams at p6 as determined by Nanostring. Colored dots represent significantly upregulated genes in offspring of X31_pre_ dams.

Each dot represents an individual pup in B-D. Statistical significance for survival curves was determined by the log-rank test (Mantel-Cox), Student’s t-test or non-parametric Student’s t-test (Mann-Whitney) in B-D.

**Figure S7: Maternal memory T cells are sufficient to protect TCR restricted offspring against heterosubtypic infection. (A)** Experimental approach to test whether maternal IAV immunity protects CD8 TCR restricted offspring against PR8 infection. **(B)** Survival of OT-I^+/-^and OT-I^-/-^littermate offspring born from naïve or X31-immune C57BL/6J dams. Pups were infected with PR8 between day 3-5 post birth. **(C)** Experimental approach to test whether maternal IAV immunity protects CD4 TCR restricted offspring against PR8 infection. **(D)** Survival of OT-II^+/-^and OT-II^-/-^offspring born from naïve or X31-immune C57BL/6J dams. Pups were infected with PR8 between day 3-5 post birth. Statistical significance for survival curves was determined by the log-rank test (Mantel-Cox)

**Figure S8: Maternal memory T cells produce IFN-y and IL-10 in the lungs of baby mice following PR8 infection. (A)** Flow cytometric confirmation of zsGreen expression by T cells in the blood of T^zsGreen^ mice.**(B)** Experimental approach (left panel) and flow cytometric analysis of maternal microchimeric T cells (zsGreen+) in the offspring lungs following IAV infection (right panel). Lungs T cells were isolated, stained with tetramer, and the tetramer-enriched fraction was analyzed by flow cytometry. T^zsG^: CD4.Cre^+/-^x zsGreen1^lslFlox/wt^; Pre-gate: CD45.2+ > Ter119- B220-CD19-Siglec-F- Ly6G- > CD90.2+. **(C)** Upper panel: Experimental approach to test whether maternal microchimeric T cells (tdTomato+) produce IFN-γ (YFP+) in the lungs of Cre-offspring following IAV infection. Lower panel: flow cytometric analysis of isolated T cells from the lungs at day 3 post-PR8 infection. T^Tom^: CD4.Cre^+/-^x tdTomato^lslFlox/wt^. Pre-gate: CD45+ > Ter119-B220- CD19- Siglec-F-Ly6G-> CD90.2+. **(D)** Upper panel: Experimental approach to test whether maternal microchimeric T cells (tdTomato+) produce IL-10 (GFP+) in the lungs of Cre-offspring following IAV infection. Lower panel: flow cytometric analysis of isolated T cells from the lungs at day 3 post-PR8 infection. T^Tom^: CD4.Cre^+/-^ x tdTomato^lslFlox/wt^. Pre-gate: CD45+ > Ter119-B220-CD19-Siglec-F-Ly6G-> CD90.2+.

**Figure S9: IL-10 deficiency on T cells does not impair recovery from primary X31 infection in adult mice and IL-10R+ cells are present in the lungs of baby mice. (A)** Experimental approach (top) and flow cytometric comparison of IL-10R expression among immune cell populations in the lungs of neonatal mice following PR8 infection. **(B)** Frequencies of immune cell populations among live CD45+ Ter119- in the IL-10R+ and IL-10R- fractions. **(C)** Weight loss assessment of adult female IL-10f/f and CD4.Cre+/WT x IL-10f/f mice following infection with 1,000 PFU X31.

**Fig S10: FluMist vaccination confers protective IAV immunity in wild-type and B cell deficient mice. (A)** Left panel: Experimental approach to test whether FluMist vaccination can protect B cell deficient mice. Right panel: Assessment of weight loss at day 4 p.i of naïve and FluMist-vaccinated μMT^+/-^and μMT^-/-^ adult littermates following lethal PR8 infection. **(B)** Representative flow plots of the lungs and spleen T cell responses of I.N or I.P vaccinated adult wild-type mice at day 10 post FluMist vaccination. Pre-gate: IV CD45-CD45.2+ > Ter119-B220-CD19-Siglec-F-Ly6G- > CD90.2+ TCRβ+ > CD4- CD8a+. **(C)** Assessment of weight loss at day 5 p.i. following lethal PR8 infection of littermate wild-type adult mice vaccinated with FluMist via the I.P or I.N routes.

