## Supplemental Figures for "Inherited maternal memory T cells protect offspring from respiratory viral infection through disease tolerance"

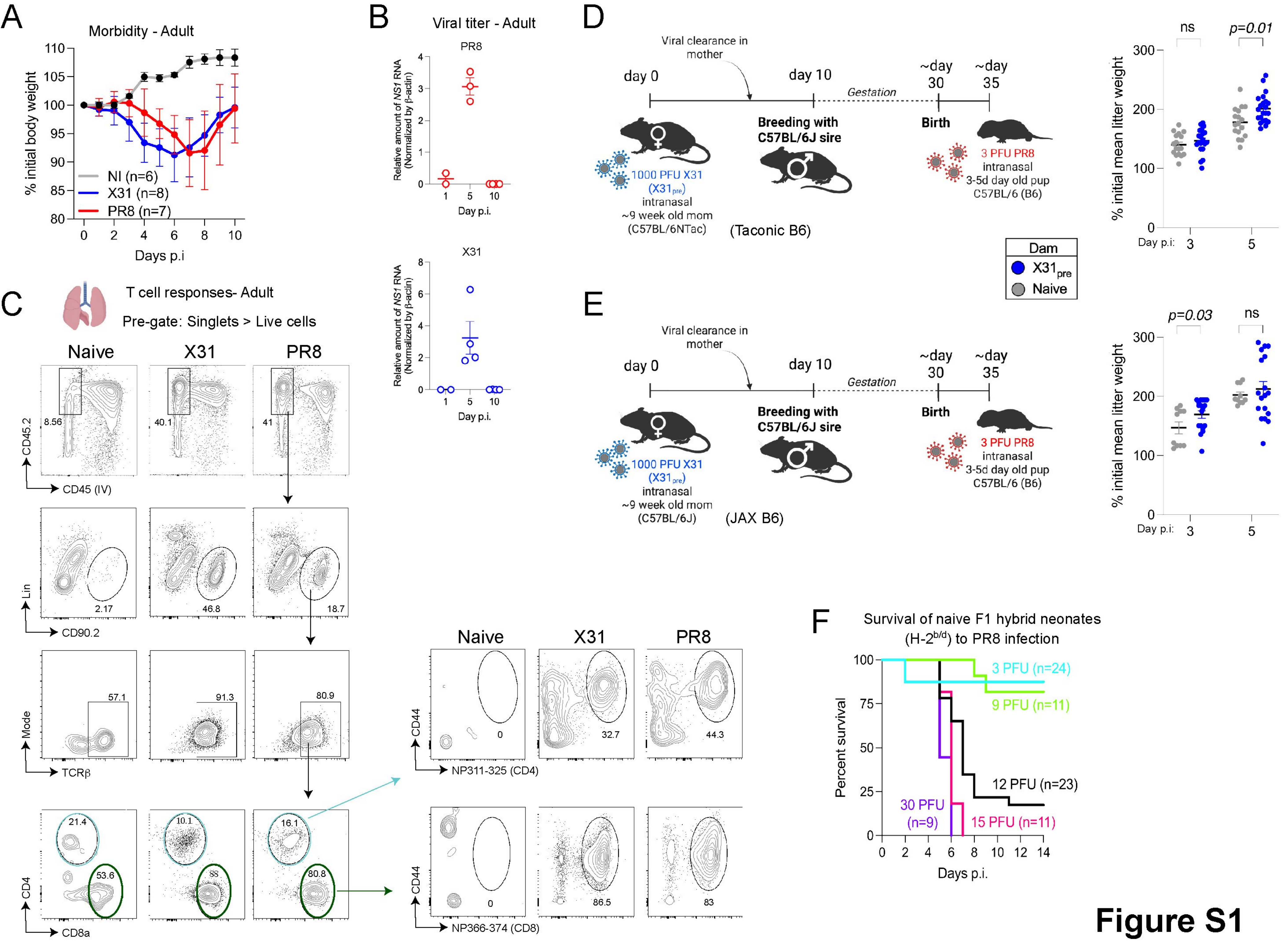


**Figure S1**
**Related to Fig1**

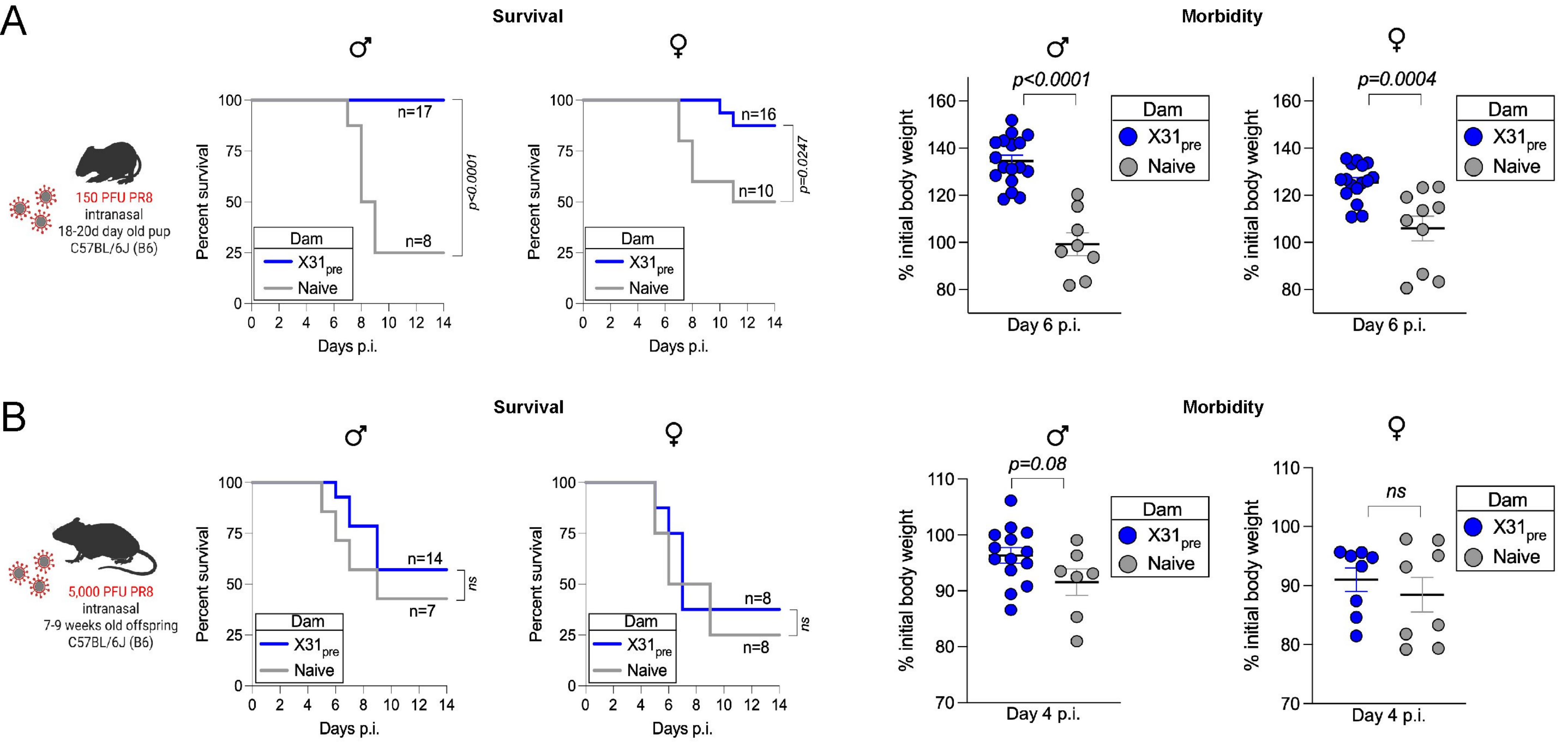


Figure S2 - Related to Fig1

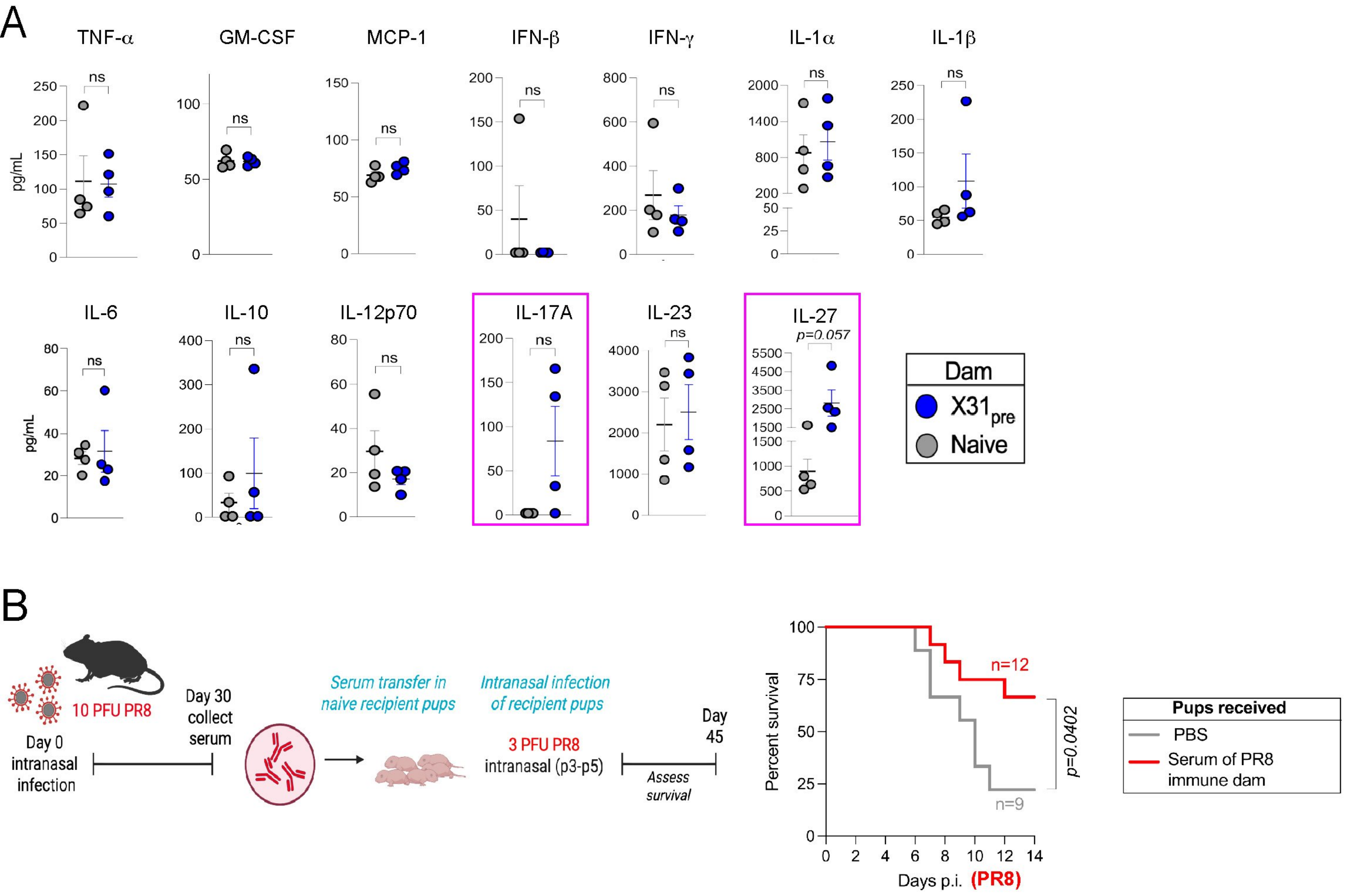


**Figure S3 - Related to Fig2**

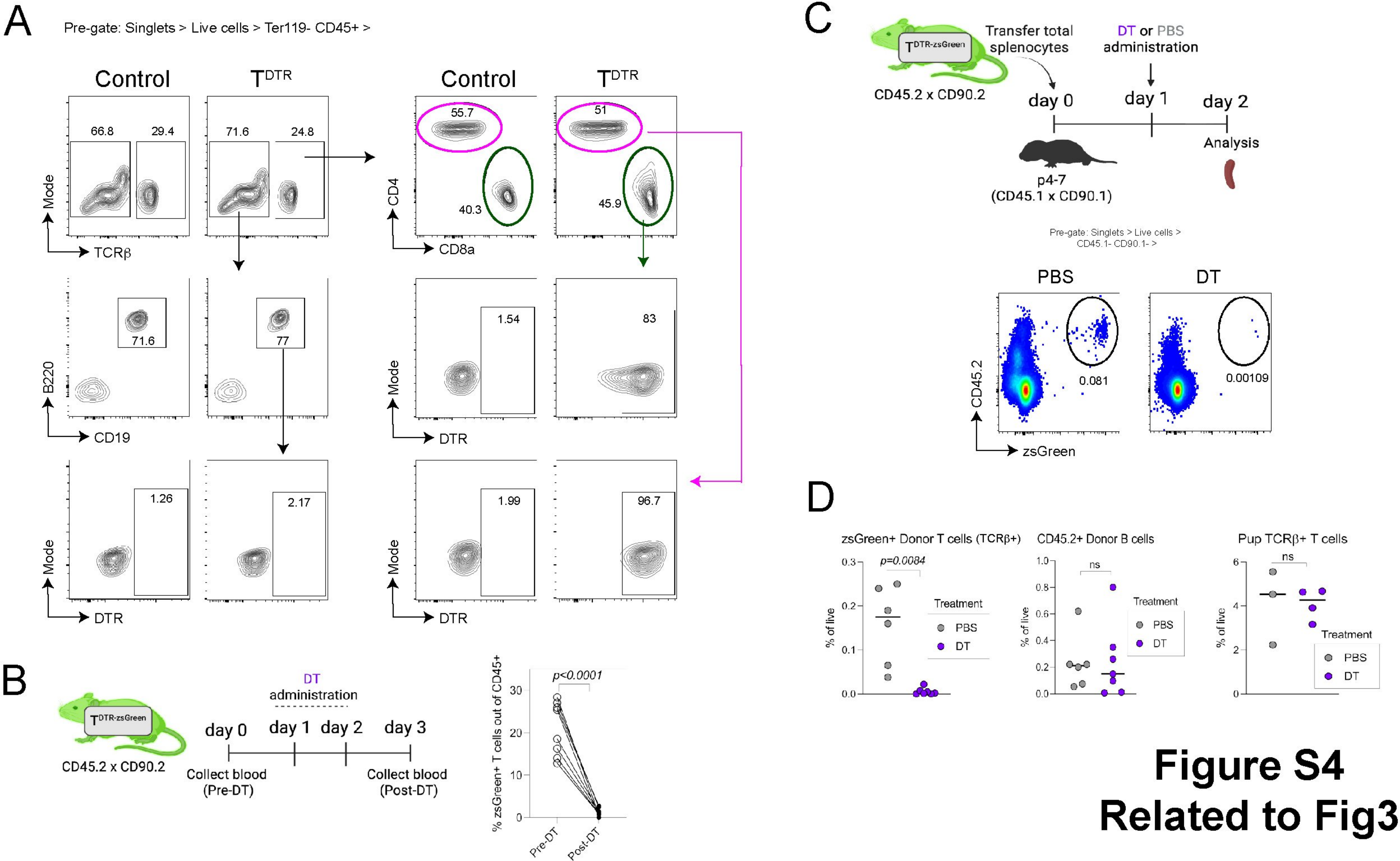


**Figure S4**
**Related to Fig3**

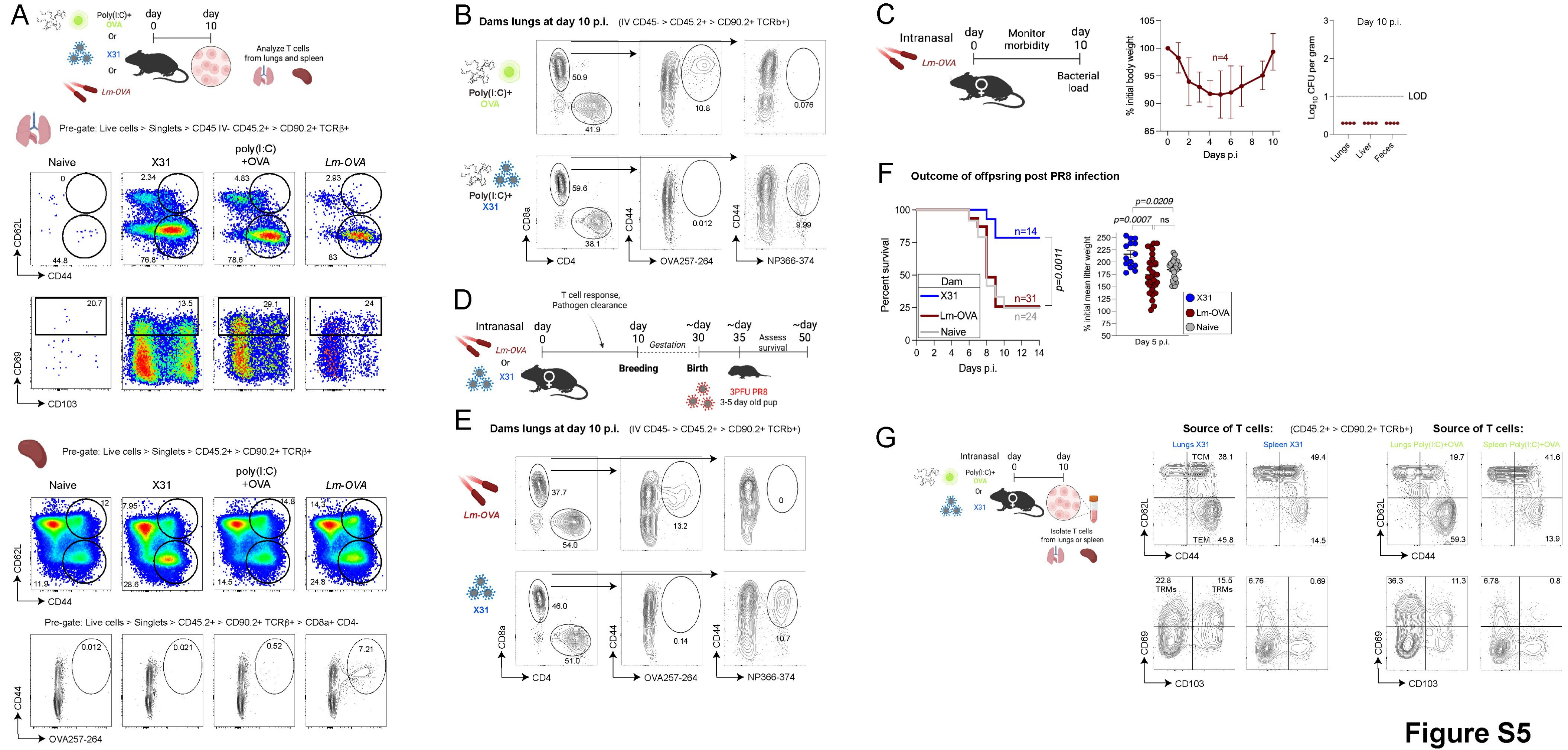


### Figure S5
Related to Fig3

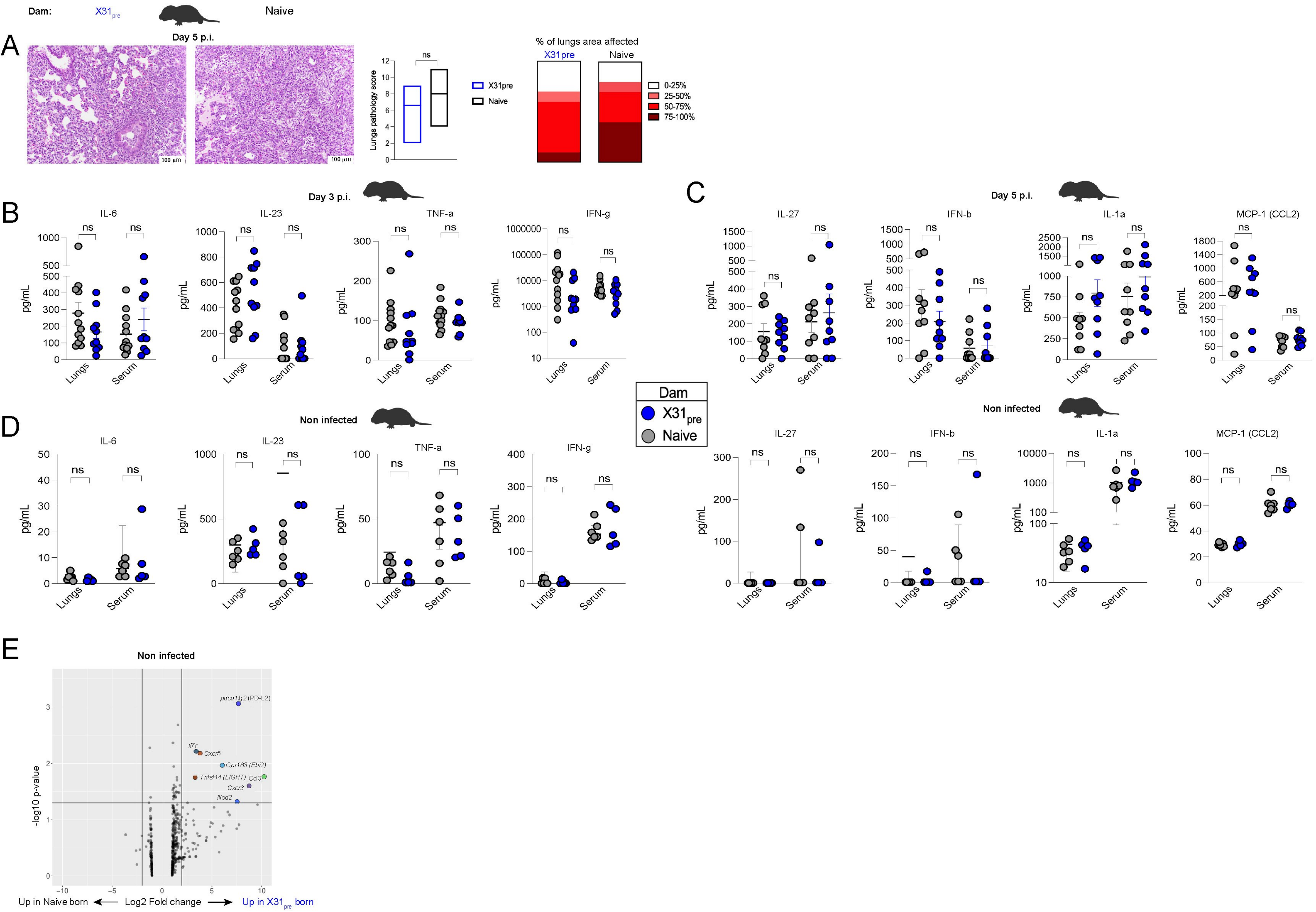


**Figure S6**
**Related to Fig4**

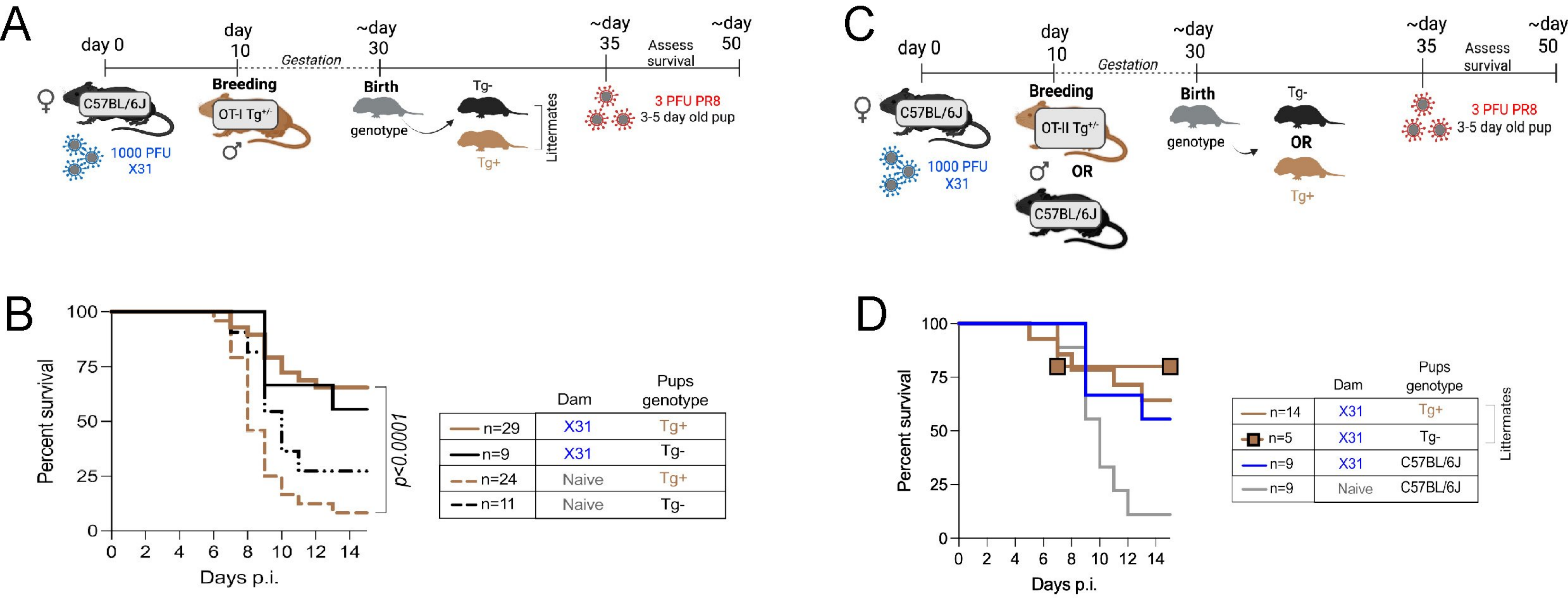


### Figure S7 - Related to Fig4

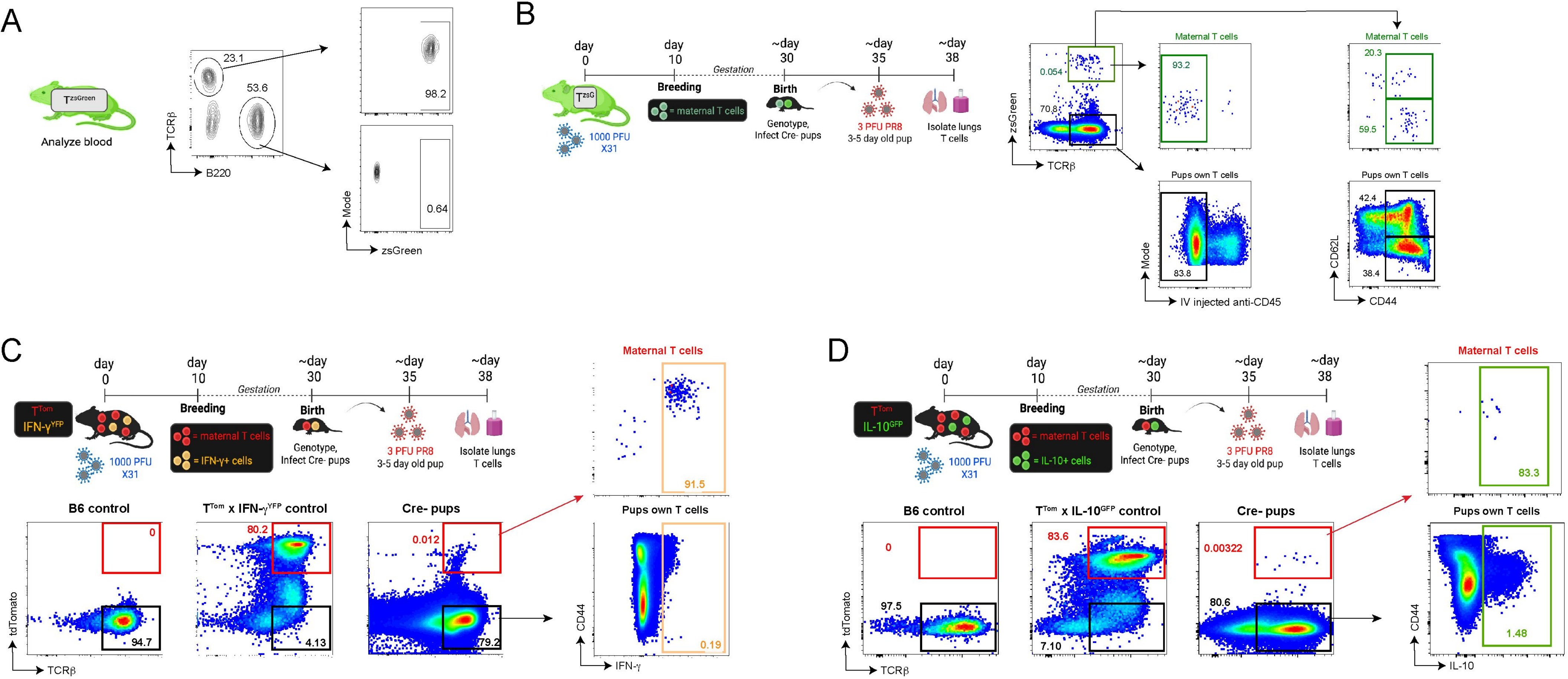


**Figure S8**
**Related to Fig5**

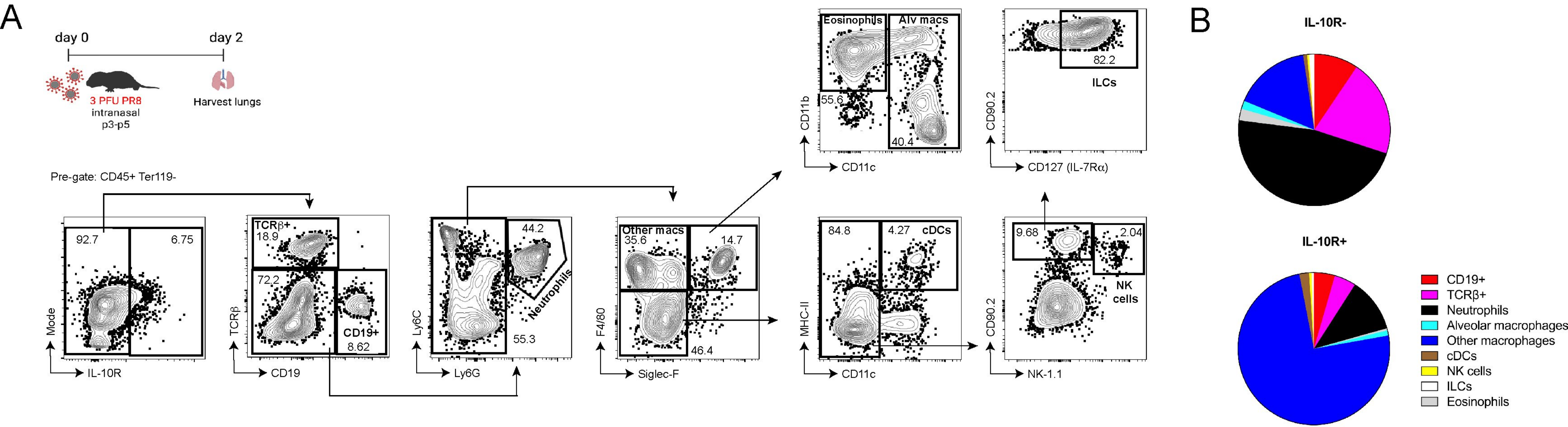


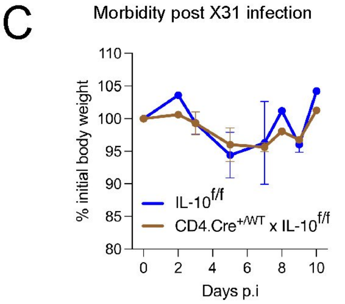


**Figure S9 - Related to Fig5**

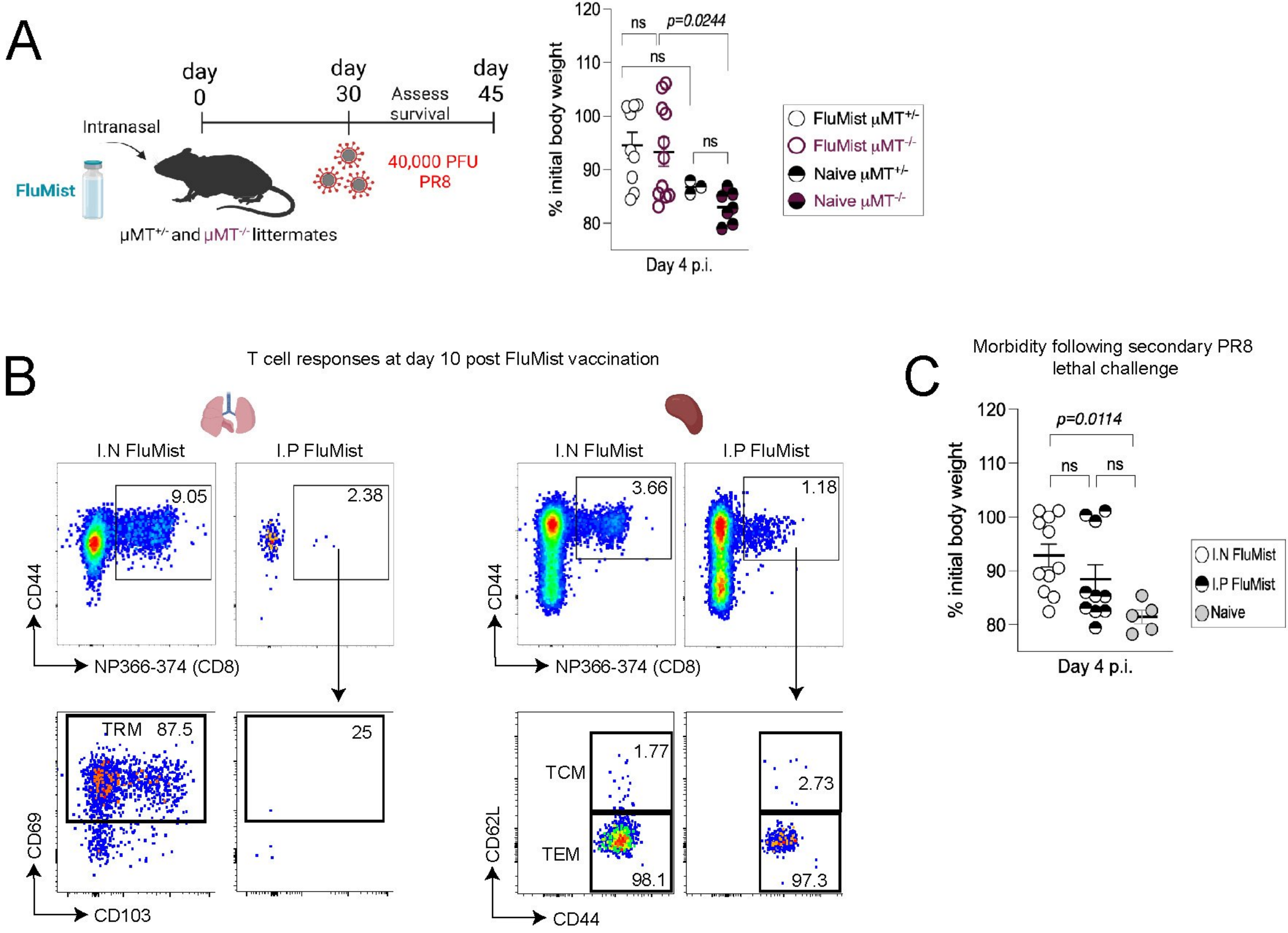


### Figure S10 - Related to Fig6
